# Mouse digit AAV gene delivery into fibroblasts regulates regenerative outcome

**DOI:** 10.1101/2025.04.22.650049

**Authors:** Vivian Jou, Scott D. Semelsberger, Jack Smerczynski, Jessica A. Lehoczky

## Abstract

The distal mouse digit tip regenerates post-amputation, while the proximal digit undergoes fibrosis. This study presents a comparative single-cell RNA sequencing-based analysis of regenerating and non-regenerating digits to computationally identify fibroblast subpopulations and genes associated with fibrosis and regeneration. To test the sufficiency of newly identified candidate genes to alter wound healing outcomes, we developed a robust adeno-associated virus gene delivery technique for digit fibroblasts. We found that overexpression of candidate pro-fibrotic genes Pcolce2 or Prelp in the blastema modifies normal regeneration and overexpression of candidate pro-regenerative factors Ccl2 or Mest in the proximal digit significantly increases bone deposition. These data demonstrate that the computational analysis combined with the AAV delivery approach presented in this study provides a powerful framework for identifying the driving factors of fibrosis and regeneration in the mammalian digit.

## INTRODUCTION

The mouse digit tip is a robust model to study innate regeneration of composite tissues in mammals^1^. Amputation through the distal phalangeal bone (P3; Figure 1A, C) results in the formation of a blastema followed by complete regeneration by 28 days post-amputation (dpa) in adult mice^2,3^. Genetic lineage analyses and single-cell transcriptomics datasets have defined the cell type heterogeneity of the mouse digit tip blastema to broadly include immune, neural, vascular, and fibroblast cells^4–7^. Fibroblasts, here defined as cells expressing Pdgfrα and Lumican, account for ∼60% of the blastema and can be computationally categorized into at least 14 distinct subtypes^6^. Previously, we identified 67 genes enriched in fibroblast subtypes that increased in population size in the blastema^6^. Of these candidate pro-regenerative genes, we further investigated Mest and determined its expression to be specific to the blastema, not generic wound-healing. With genetic loss of function experiments, we defined its role in the immune system to mediate proper bone regeneration^8^. However, the sufficiency of Mest, and the other candidate genes, to induce regeneration remains unknown.

**Figure 1.**
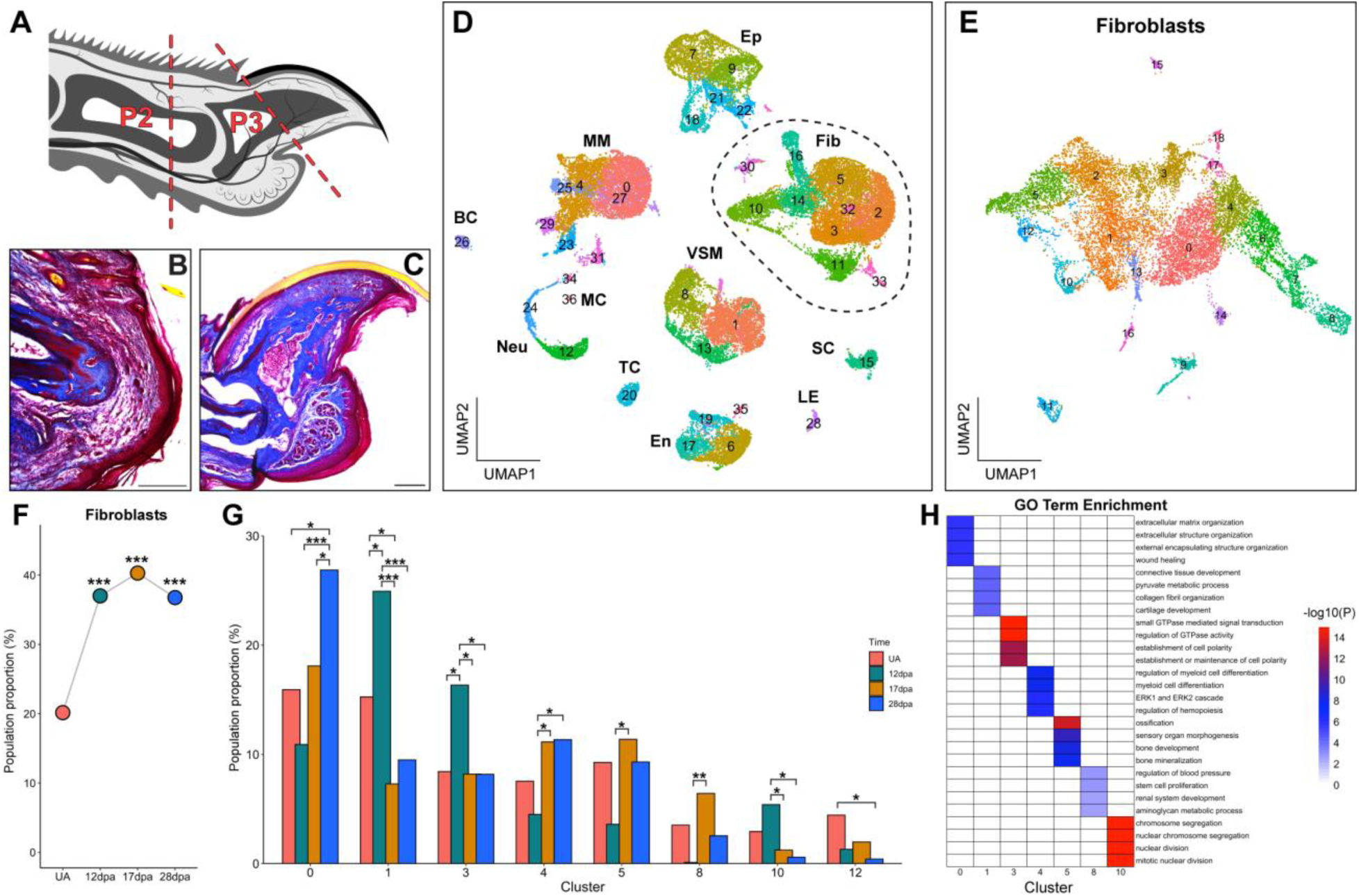
Fibroblast heterogeneity and dynamics during proximal P2 digit fibrosis. (A) Schematic cross section of mouse digit; dashed lines indicating amputation planes through P2 and P3 digit bones. Masson’s trichrome histology of 28dpa (B) P2 fibrosis and (C) P3 regeneration. (D) UMAP plot of integrated P2 UA, 12dpa, 17dpa, and 28dpa scRNAseq Seurat defined clusters. BC: B cells, En: endothelial cells; Ep: epithelial cells, Fib: fibroblasts, LE: lymphatic endothelial cells, MC: mast cells, MM: macrophages/monocytes, Neu: neutrophils, SC: Schwann cells, TC: T cells, VSM: vascular smooth muscle. Dashed circle highlights fibroblasts that subsetted and reclustered in (E). Differential proportion analysis of (F) proportion of total fibroblasts in the UA, 12dpa, 17dpa, and 28dpa P2 digit and (G) fibroblast subtypes with significant population size changes during fibrosis. (H) Top four GO terms enriched for each fibroblast subtype in (G). Scale bars: 250µm; *p<0.05, **p<0.01, ***p<0.001.

In contrast to the P3 digit tip, amputations through the proximal digit, such as mid-digitally through the second phalangeal bone (P2; Figure 1A-C), do not regenerate and instead heal via fibrosis^9–11^. Thus, the P2 digit amputation model is ideally suited for comparing regenerative and fibrotic fibroblasts and for regenerative sufficiency experiments^11,12^. To this point, innate differences between P2 and P3 fibroblasts have been noted, though these studies do not address fibroblast heterogeneity or subtype specific gene expression and function^7,13^. A comprehensive comparative analysis of fibroblast subtypes, their gene expression, putative functions, and population dynamics in P3 regeneration versus P2 fibrosis remains to be done. Separately, the P2 digit amputation is also an ideally suited platform to functionally test the sufficiency of candidate genes to induce regeneration in innately non-regenerative tissues. Indeed, previous studies have introduced exogenous factors including iPSCs, proteins, and ECM components into P2 post-amputation wounds using techniques including conditional genetics, bead implantation, direct injection, or systemic delivery^14–19^. While these reports collectively suggest that full regeneration can be induced from a P2 amputation given the correct combination of exogenous factors, the currently utilized techniques lack the flexibility and scalability needed for high-throughput screening of candidate genes.

In this paper, we present single-cell RNA sequencing (scRNAseq) time courses for stage-matched mouse P2 and P3 post-amputation digits and directly compare fibroblast subtypes throughout digit fibrosis and digit tip regeneration. We find that P2 and P3 post-amputation digits are enriched for distinct fibroblast subtypes, indicating innate cell type differences between digit fibrosis and regeneration. We then identified 98 candidate pro-fibrotic and 27 candidate pro-regenerative genes. Towards high throughput functional assessment of these candidate genes to drive fibrosis or regeneration, we developed fibroblast-specific adeno-associated virus (AAV) mediated gene delivery for local, transient gene expression. As proof of concept, we evaluated three candidate pro-fibrotic genes for sufficiency to inhibit regeneration following P3 amputations and three candidate pro-regenerative genes for sufficiency to induce regeneration following P2 amputations. We demonstrate that AAV delivery of candidate pro-fibrotic ECM regulator genes Pcolce2 or Prelp into blastemal fibroblasts alters 3D bone morphology during regeneration, and that AAV delivery of blastemal genes Ccl2 or Mest into P2 fibroblasts induces bone regeneration in the non-regenerative digits.

## RESULTS

### Distinct fibroblast subtype heterogeneity in P2 fibrosis vs P3 regeneration yields candidate genes for functional testing

To assess fibroblast heterogeneity during fibrosis of the P2 post-amputation digit, we performed scRNAseq of unamputated (UA) and 12-, 17-, and 28-days post-amputation (dpa) P2 wildtype FVB mouse digits. The genetic background and timepoints were matched to our previously reported P3 digit tip blastema dataset (GSE143888) to facilitate direct comparison of fibroblast subtypes^6^. Following sequencing, quality control, and filtering, 11834 (UA), 7208 (12dpa), 8268 (17dpa), and 11752 (28dpa) P2 cells were used for analysis (Figure S1A-D). Canonical markers were used to assign broad cluster identities (Figure S1E), revealing cell-type heterogeneity at all time points, consisting of fibroblast, immune, vascular smooth muscle (VSM), endothelial, and epithelial cells (Figure S1A-D). Epithelial cells were underrepresented due to minimized epidermis collection during digit dissection. Integration of UA, 12-, 17-, and 28dpa scRNAseq data reinforced the cell type heterogeneity found in the homeostatic and fibrosing P2 digit (Figure 1D), and while distinct in genetic background, stage and cell type proportions, our results were consistent with GSE135985^7^ (Figure S1F).

To specifically address fibroblast heterogeneity, we subsetted and reclustered all P2 fibroblasts, which resulted in 19 computationally defined clusters, indicative of subpopulations as was found in the P3 digit blastema (Figure 1E). Consistent with the P2 digit undergoing fibrosis, the total proportion of fibroblasts significantly increased at all post-amputation stages as compared to the unamputated digit (Figure 1F). To identify which fibroblast subtypes predominate at various stages, we assessed their population proportions over time. Differential proportion analysis revealed that clusters 0, 1, 3, 4, 5, 8, 10, and 12 were the only clusters that exhibited significant population changes during fibrosis (Figure 1E, G). While each cluster had a distinct population dynamics profile, they broadly fell into three categories. 1) The fibroblast clusters 0, 4, 5, and 8 decreased in proportion at 12dpa then increased by 17dpa or 28dpa. Analysis of cluster-specific differentially expressed genes (DEGs), GO terms, and KEGG pathway enrichment suggests clusters 0, 4, 5, and 8 are associated with ECM organization, myeloid cell differentiation, ossification, and regulation of hematopoiesis, respectively (Table S1, Figure 1G, H, S1G). 2) The fibroblast clusters 1, 3, and 10 increased in proportion at 12dpa then became depleted at 17dpa and 28dpa; these populations are predicted to be involved in collagen organization, Rap1 signaling, and mitosis, respectively (Table S1, Figure 1G, H, S1G). 3) Cluster 12, which is associated with tissue development, was the only population to be depleted at all post-amputation time points (Table S1, Figure 1G, S1G, H). We verified that the top DEGs associated with each significant cluster were also differentially expressed across time; for example, Fbn2 is a significant DEG at 12dpa for cells in clusters 1 and 3, while Pcolce2 is a significant DEG at 28dpa for cells in clusters 0 and 4 (Figure S1I, J, Table S2). These results suggest that, similar to digit tip regeneration, different fibroblast subpopulations with distinct biological functions contribute to varying stages of wound healing.

To determine how fibroblast subtypes correlate between P2 fibrosis and P3 regeneration, we integrated our stage matched P2 and P3 scRNAseq datasets (UA, 12-, 17-, and 28dpa, Figure S2A). Common cell-type heterogeneity was found between the P2 and P3 tissues; however, the proportions of these populations varied significantly (Figure S2B). Notably, total P3 fibroblasts were 3.1x, 2.1x, 1.5x, and 2x more populous as compared to all P2 fibroblasts at homeostasis, 12dpa, 17dpa, and 28dpa, respectively (Figure S2A’-D’). Fibroblasts at each matched time point from the P2-P3 integrated dataset were subsetted and reclustered for comparative analysis of subpopulations (Figure 2A-D). While P2 and P3 derived cells were found in all clusters, the mixing was strikingly non-homogeneous (Figure 2A’-D’). Differential proportion analysis revealed that there were 10 UA, 3 12dpa, 7 17dpa, and 10 28dpa fibroblast subpopulations significantly skewed in proportion between P2 and P3 tissues (Figure 2E-H). This indicates that P2 and P3 fibroblasts are quite distinct, even in homeostasis, corroborating previous studies^7,13^.

**Figure 2.**
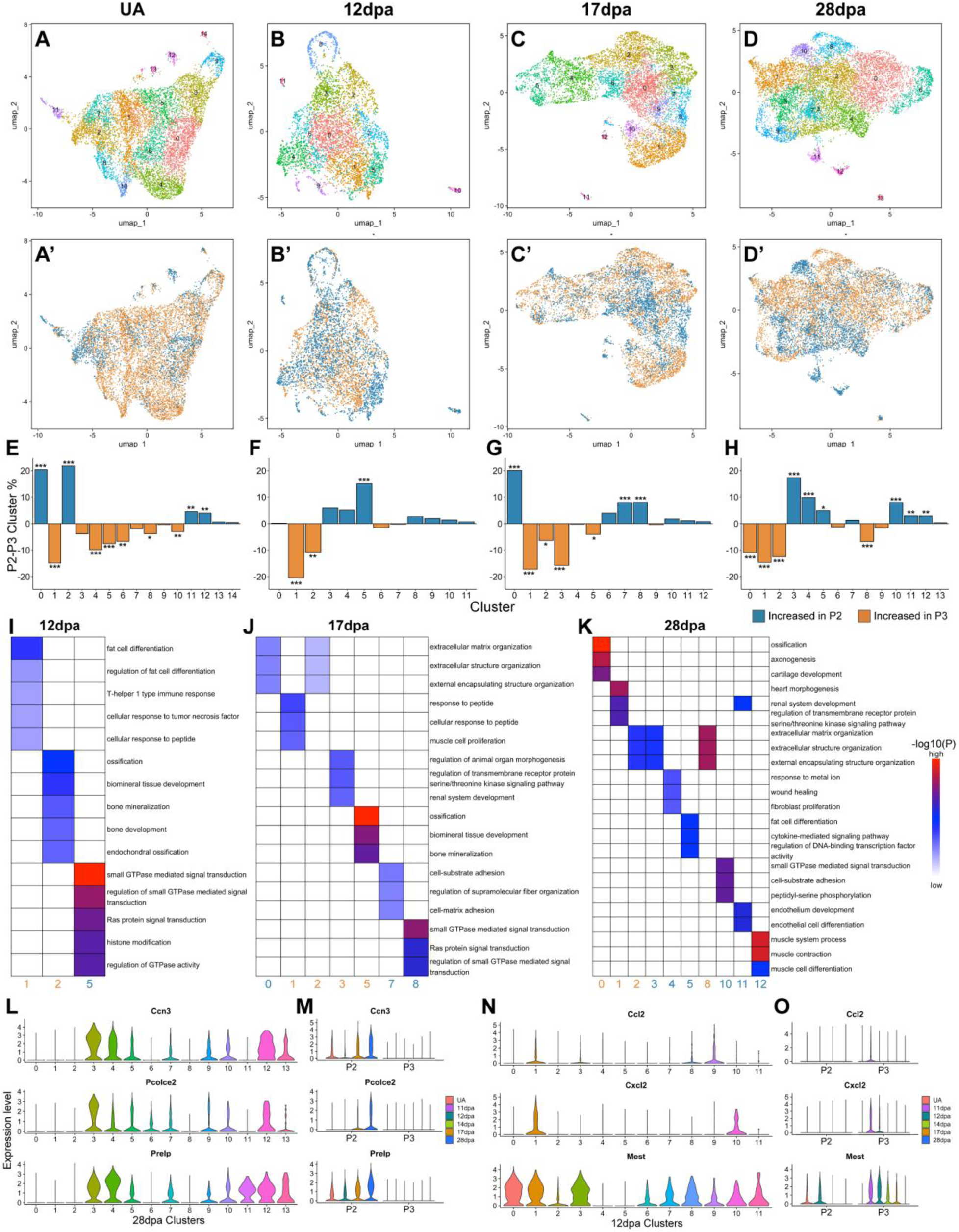
Computational analysis of fibroblasts through P2 fibrosis compared to P3 regeneration. (A-D) UMAP plots of Seurat integrated and clustered P2 and P3 fibroblasts at (A) UA, (B) 12dpa, (C) 17dpa, and (D) 28dpa; (A’-D’) UMAP plots from (A-D) colored by P2 (blue) or P3 (orange) digit origin. (E-H) Differential proportion analysis of fibroblast subtype population skewing between P2 (blue) and P3 (orange) in UA, 12dpa, 17dpa, and 28dpa digits. (I-K) Top five enriched GO terms for significantly skewed P2 or P3 fibroblast subtypes from (F-H) at (I) 12dpa, (J) 17dpa, and (K) 28dpa. (L-M) Examples of candidate pro-fibrotic genes; violin expression plots of P2 enriched genes Ccn3, Pcolce2, and Prelp in (L) 28dpa P2-P3 integrated fibroblasts subtypes (from D and H) and in (M) total fibroblasts through P2 fibrosis and P3 regeneration. (N-O) Examples of candidate pro-regenerative genes; violin expression plots of P3 enriched genes Ccl2, Cxcl2, and Mest in (N) 12dpa P2-P3 integrated fibroblast subtypes (from B and F) and in (O) total fibroblasts through P2 fibrosis and P3 regeneration. *p<0.05, **p<0.01, ***p<0.001.

To assign putative function to these P2 or P3 enriched subpopulations we performed GO term and KEGG pathway enrichment (Figure 2I-K, Figure S2E-G). Broadly over all post-amputation stages, P3 enriched fibroblast subtypes had gene signatures of bone development and ossification, while P2 enriched fibroblast subtypes involved GTPase mediated signaling and cell-adhesion (Figure 2I-K). In addition, upregulated pathways in immune response, such as TNF and IL-17 signaling, were associated with 12dpa and 17dpa P3 enriched fibroblast subtypes, but at 28dpa this biological profile is associated with the P2 fibroblast subtype in cluster 5 (Figure S2E-G). This suggests prolonged inflammation in the P2 wound, a canonical characteristic of the fibrotic response in many organs^20,21^. Separately, ECM organization GO terms were found for both P2 and P3 fibroblast subpopulations at 17dpa and 28dpa. Interestingly, the genes significantly associated with this GO term differed between P2 and P3 fibroblasts, suggesting distinct ECM environments between regeneration and fibrosis (Figure S2H).

For downstream in vivo analysis, we sought to define candidate pro-fibrotic genes associated with P2-enriched fibroblast subtypes and candidate pro-regenerative genes associated with P3-enriched fibroblast subtypes. All P2 and P3 fibroblasts, including those from our previously reported 11dpa and 14dpa P3 blastema dataset were computationally integrated and clustered^6^ (Figure S2I, J). To identify candidate pro-fibrotic genes, we isolated the DEGs from 28dpa significantly enriched P2 clusters (28dpa clusters 3, 4, 5, 10, 11, and 12; Figure 2H, Table S3) with a minimum percentage of cells greater than 0.25, an average log2 fold-change greater than 0.58, and an adjusted p-value less than 0.05. These genes were filtered for those highly upregulated in 28dpa P2 fibroblasts as compared to UA, 12dpa, and 17dpa P2 fibroblasts (Table S2). Of the resulting 98 candidate pro-fibrotic genes (Table S4), we selected Ccn3 (cellular communication network factor 3), Pcolce2 (procollagen C-endopeptidase enhancer 2), and Prelp (prolargin; proline/arginine rich end leucine rich repeat protein) for further assessment (Figure 2L, M). Separately, to identify candidate pro-regenerative genes, we isolated the DEGs from 12dpa significantly enriched P3 clusters (12dpa clusters 1 and 2; Figure 2F, Table S5) with a minimum percentage of cells greater than 0.25, an average log2 fold-change greater than 0.58, and an adjusted p-value less than 0.05. These genes were further filtered based on our previously identified blastema enriched genes and resulted in 27 candidate pro-regenerative genes (Table S6). Ccl2 (C-C motif chemokine ligand 2), Cxcl2 (C-X-C motif chemokine ligand 2), and Mest (mesoderm specific transcript) were selected for functional testing^6^. Ultimately, these two lists of genes represent factors that may be integral in determining whether the tissue heals via fibrosis or regeneration.

### AAV2/6 injection yields robust transient gene expression in digit fibroblasts

Our comparative analysis of P2 and P3 fibroblast subtypes identified numerous putative pro-fibrotic and pro-regenerative candidate genes (Tables S4, S6). Ideally, every candidate gene would be evaluated in vivo, however, existing techniques for gene or protein modulation in mouse digits are limited by reagents and throughput. In turn, we sought to develop a flexible, higher-throughput, gene candidate screening platform for the digit, and turned to adeno-associated viruses (AAV). In recent years, AAVs have become a popular tool for gene delivery due to their non-pathogenicity, flexibility in cell type tropism, and scalability for large studies^22–25^. We began by assessing five pseudotypes of AAV-CAG-GFP: 6, 8, 9, 7m8, and PHP.S, chosen based on previous studies supporting infectivity in mesenchymal cells^25–28^. Five cohorts of wildtype adult mice underwent distal digit tip amputations (P3 regeneration; Figure 1A) while separate cohorts underwent mid-digital amputations (P2 fibrosis; Figure 1A). At 6dpa, 2uL of high titer AAV was injected locally via Hamilton syringe, using one serotype per P2 or P3 cohort (Figure S3A). All digits were analyzed for GFP expression at 12dpa (6 days post injection (dpi)), and results were consistent between P2 and P3 tissues. While minimal GFP expression was found following AAV8 or 9 infections, significant expression was found in the AAV6, 7m8, and PHP.S injected digits (Figure 3A-D). AAV6 infection resulted in 11.2x and 4.0x increases in GFP expression than AAV8 in the P2 and P3 digits, respectively, and 271.6x and 4.8x increases versus AAV9 in the P2 and P3 digits, respectively (Figure 3A-D). Further analysis revealed that AAV6 infection remains local to the digit injection site (Figure S3B, C) and infects fibroblasts including VIM^+^ (Figure S3D-D”) and RUNX2^+^ cells (Figure S3E-E”). No GFP expression colocalizes with the epidermis or F4/80^+^ macrophages (Figure S3F-G’). To move forward with our experiments, we selected AAV6 for its high infectivity of mesenchymal cells in both P3 and P2 post-amputation digits.

**Figure 3.**
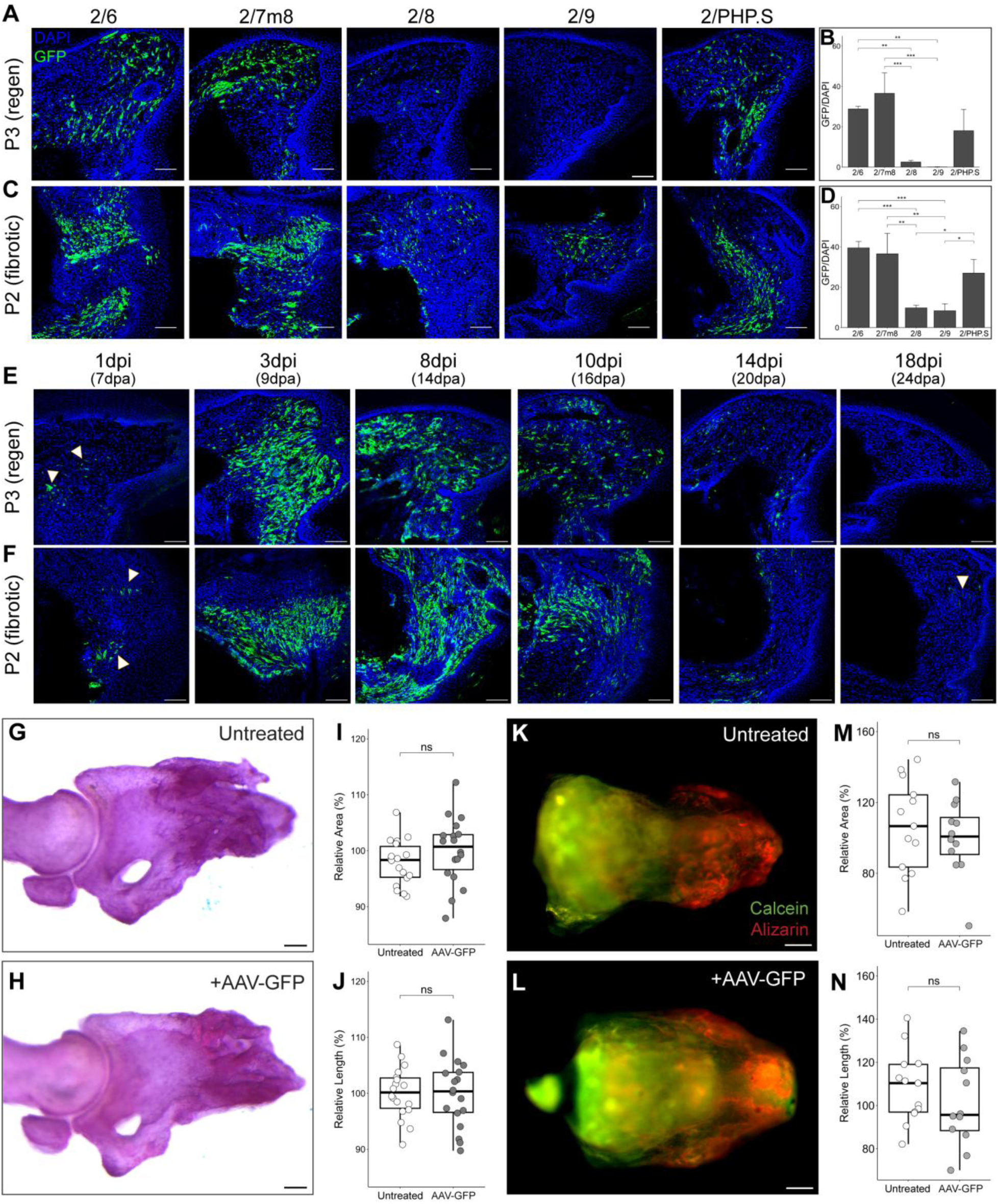
AAV2/6 infects mouse P2 and P3 post-amputation fibroblasts. Anti-GFP section IHC of 12dpa/6dpi (A) P3 regenerating digits and (C) P2 fibrosing digits following AAV-CAG-GFP injection with pseudotypes 2/6, 2/7m8, 2/8, 2/9, or 2/PHP.S. (B, D) Image quantification of GFP signal across AAV pseudotypes for (B) P3 and (D) P2 12dpa/6dpi digits. (E-F) Anti-GFP section IHC of 1-, 3-, 8-, 10-, 14- and 18dpi (E) P3 and (F) P2 digits following AAV2/6-CAG-GFP injection; white arrowheads = examples of infected cells. (G, H) Representative alizarin red skeletal stained P3 28dpa regenerated bones following (G) no AAV injection (untreated) and (H) AAV-GFP injection. 2D quantification of P3 bone (I) area and (J) length. (K, L) Representative calcein (green) and alizarin complexone (red) differentially stained P2 28dpa bones for (K) untreated and (L) AAV-GFP injected digits. 2D quantification of new bone (red) (M) area and (N) length. Scale bars = 100µm. ns = not significant, *p<0.05, **p<0.01, ***p<0.001. Data are shown as median ± interquartile range.

To determine the post-injection timing of AAV6-CAG-GFP expression, we assessed GFP expression at additional time points of 1, 3, 8, 10, 14, and 18dpi (7, 9, 14, 16, 20, and 24dpa, respectively) (Figure 3E, F, S3H, I). For both P2 and P3 post-amputation digits, several cells adjacent to the injection site express GFP as early as 1dpi (Figure 3E, F). In P3 regenerating digits, there is an 8.7x increase in GFP/DAPI signal at 3dpi (Figure 3E, S3H; 1 vs 3dpi p = 2.1E-7) and this expression persists until 10dpi, when GFP expression decreases by 3.6x from 8dpi (Figure 3E, Figure S3H; 8 vs 10dpi p=7.1E-5). In P2 fibrosing digits, there is a 5.5x increase from 1dpi to 3dpi (1 vs 3dpi p=4.7E-6) and peak GFP signal is found at 6dpi (Figure 3F, S3I). Expression then significantly decreases at each subsequent time point, with the most significant drop in GFP signal occurring between 8 and 10dpi (1.9x decrease, p=9.8E-4). By 14dpi, GFP expression is restricted to only a few sparse cells (Figure 3F, S3I). Thus, in both P2 and P3 digits, the maximum overexpression window is from 3-10dpi.

To assure no AAV-related phenotypes were found, we compared infected and untreated P2 and P3 digits. To assess P3 bone regeneration, untreated and AAV-GFP infected P3 digit tips were analyzed at 28dpa (22dpi) by quantification of alizarin red skeletal staining (Figure 3G, H). Importantly, no significant differences were found in area or length of the P3 bone, indicating that AAV infection itself does not perturb regeneration (Figure 3I, J). To determine if there was any AAV infection related change in bone repair between untreated and AAV-GFP infected P2 digits, we analyzed differential calcein and alizarin complexone staining (Figure 3K, L). Unlike the P3 digit tip, the absence of reliable P2 anatomical landmarks results in subtle amputation position variation, thus these vital bone dyes can clearly delineate pre-amputation (green) and post-amputation (red) bone. Quantification of area and length of new bone growth revealed no significant differences between untreated and AAV-GFP infected 28dpa P2 digits (Figure 3M, N). These results demonstrate AAV6 is a robust and efficient tool to use for local and transient gene expression manipulation in both P2 and P3 post-amputation digit fibroblasts.

### AAV delivery of P2-derived candidate pro-fibrotic genes to P3 blastema alters regeneration

To determine if gene expression specific to P2 enriched fibroblast subtypes function in driving fibrosis and/or repressing regeneration, we aimed to leverage this AAV delivery platform to screen our computationally identified candidates (Tables S4, S6). As proof of principle, we tested three of our computationally identified candidate pro-fibrotic genes: Ccn3, Pcolce2, and Prelp (Table S4). As described above, these were chosen for their distinct expression in significantly enriched fibrotic subclusters (Figure 2L, M). Additionally, all three are matrisome associated genes with distinct functions in fibrosis: Ccn3 is a glycoprotein that has been shown to both promote and inhibit fibrosis in various tissue models, Pcolce2 is a procollagen processing enzyme found to promote cardiac fibrosis, and Prelp is a member of the small leucine-rich proteoglycan family and found enriched in heart, liver, and adipose tissue fibrosis^29–38^. High titer AAV6 was produced for mouse Ccn3, Pcolce2 and Prelp cDNA constructs. Mice underwent distal P3 amputations and at 6dpa, each experimental digit was injected with 2µL of a single recombinant AAV. Eighteen infected digits from each cohort were collected at 6dpi to validate AAV-derived expression by qPCR analysis. Compared to AAV-GFP infected control digits, we observed 9624x, 157x, and 171x increases in Ccn3, Pcolce2, and Prelp expression, respectively (Figure S4A-C).

The digits from remaining mice of all cohorts were analyzed at 28dpa (22dpi) for any regenerative phenotypes. Masson’s trichrome histology reveals that all digits in each treatment group regenerated with no gross tissue composition differences (Figure S4D-G). Additional digits were assessed by alizarin red skeletal staining for P3 2D bone area and length quantification (Figure 4A-D). AAV-Ccn3 infected digits exhibited no significant change in either bone area or length as compared to AAV-GFP digits (Figure 4B, E, F). In contrast, 28dpa AAV-Prelp infected digits were 7% shorter in bone length as compared to AAV-GFP infected digits (Figure 4D-F; p=0.036), consistent with inhibiting regeneration. Surprisingly, 28dpa AAV-Pcolce2 infected digits had 11% increased bone area and 10% increased bone length as compared to AAV-GFP infected digits (Figures 4C, E, F; p=9.10E-05 and 2.90E-04, respectively).

**Figure 4.**
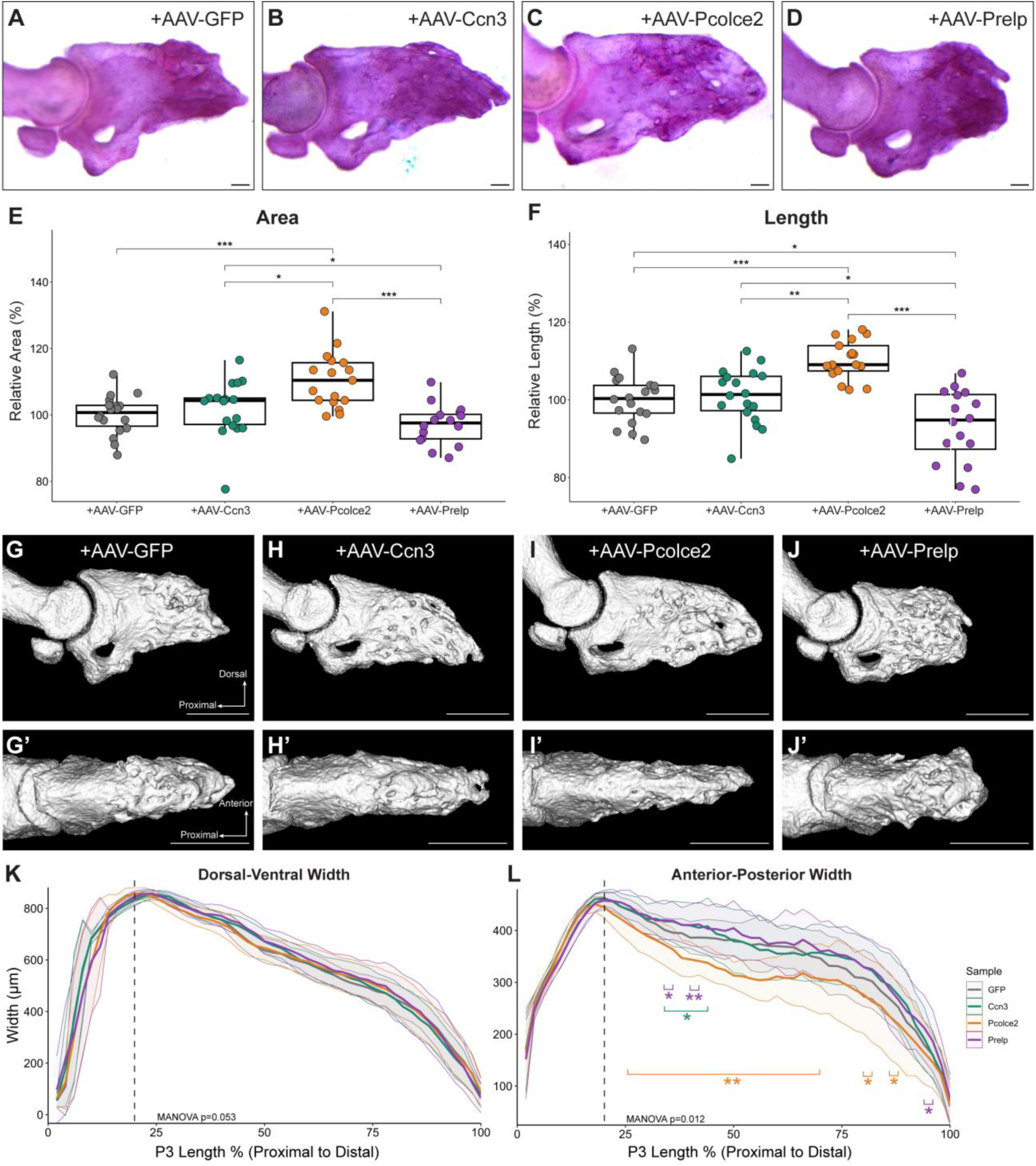
AAV delivery of candidate pro-fibrotic genes to P3 regenerating digits. Representative alizarin red skeletal stained 28dpa P3 bones following AAV infection of (A) GFP, (B) Ccn3, (C) Pcolce2, and (D) Prelp; scale bars = 100µm. Quantification of P3 2D bone (E) area and (F) length relative to AAV-GFP infected digits. Representative (G-H) lateral and (G’-H’) dorsal view microCT images of 28dpa AAV infected digits tips; scale bars = 500µm. (K, L) MicroCT-derived (K) dorsal-ventral or (L) anterior-posterior width measurements versus length for AAV infected cohorts. Thick colored lines report average for each cohort; adjacent colored shaded areas report standard deviation. Only statistical comparisons against GFP samples are shown. ns = not significant, *p<0.05, **p<0.01, ***p<0.001.

To further determine any phenotypic changes in the 3D structure of these digits, we performed microCT analysis on these bones (Figure 4G-J). No significant differences were found in volume or surface area of the 28dpa P3 bones for any of AAV-Ccn3, Pcolce2, or Prelp cohorts as compared to the AAV-GFP digits (Figure S4H, I). These data were counterintuitive given the 2D bone area and length findings for AAV-Pcolce2 and AAV-Prelp infected digits (Figure 3E-F). Thus, we reasoned differences along the anterior-posterior (A-P) axis could account for unchanged volumes and surface areas. Indeed, while the dorsal-ventral (D-V) measurements were consistent across treatment groups (Figure 4G-K; MANOVA p = 0.0534), the A-P bone widths significantly differed with the overexpressed gene (Figure 4G’-J’, L; MANOVA p= 0.012). Notably, as compared to AAV-GFP infected digits, AAV-Pcolce2 bones were significantly thinner while AAV-Prelp bones were thicker throughout the P-D bone length (Figure 4L). These A-P data reconcile the 2D bone length and 3D bone volume and surface area findings. Thus, while none of these candidate pro-fibrotic genes were individually sufficient to fully inhibit regeneration, Prelp and Pcolce2 overexpression affected the 3D structure of the regenerated bone, supporting a role in collagen remodeling and bone deposition.

### P2 bone regeneration is induced by AAV delivery of candidate pro-regenerative genes

To identify if genes specific to the enriched P3 blastemal fibroblast subtypes function to stimulate regeneration and/or suppress fibrosis, we used AAV infection in non-regenerating P2 amputations. As proof of principle, we tested three genes from our computationally identified pro-regenerative candidates (Figure 2N, O, Table S6): Ccl2, Cxcl2, and Mest. These genes were selected for their high enrichment in blastemal specific clusters (Table S5, S6) as described above. Furthermore, Ccl2 specifically recruits macrophages and was found to promote regeneration in tissues such as the heart, articular cartilage, and bone; Cxcl2 recruits neutrophils and stimulates liver regeneration; Mest is necessary for proper immune cell recruitment and clearance to support digit tip regeneration^8,39–43^. Mouse Ccl2, Cxcl2, and Mest cDNAs were cloned and used to produce high titer AAV6 for injection. Four cohorts of adult wildtype mice underwent P2 digit amputations. At 6dpa, 2uL of AAV was injected into the healing digit, one gene per mouse/cohort. Eighteen digits from each cohort were collected at 6dpi (12dpa) to verify AAV-derived gene expression with qPCR and revealed a 23.4x, 5.7x, and 3.4x increase of Ccl2, Cxcl2, and Mest expression as compared to AAV-GFP infected digits, respectively (Figure S5A-C).

To characterize the sufficiency of each gene to promote regeneration, 28dpa (22dpi) P2 bones were analyzed following Ccl2, Cxcl2, or Mest AAV infection. Masson’s trichrome staining revealed that AAV-Mest infected digits had an aggregation of connective tissue abutting the distal repaired bone, that was significantly larger and unlike the distal tissue found in any of the other cohorts (Figure 5A-E). While Safranin O/Fast Green staining indicated these cells were not chondrocytes (Figure S5D), some of these cells expressed the osteoblast marker Sp7, beyond what was found in the AAV-GFP infected digits (Figure S5E, F). This indicates an enrichment of osteolineage cells in the AAV-Mest digits, thus, we aimed to quantify the amount of bone repair utilizing differential calcein and alizarin complexone fluorescence (Figure 5F-I). AAV-Ccl2 infected digits averaged a 36% increased area and 12% increased length for new bone as compared to the AAV-GFP injected digits; however, these findings were not statistically significant perhaps due to the interdigit variation (Figures 5G, J, K). Consistent with our hypothesis and prior reports supporting Mest as a pro-regenerative gene^6,8^, AAV-Mest infected digits exhibited a 65% increased area and 25% increased length of P2 bones compared to AAV-GFP controls (Figure I-K; p=4.8E-4 and 2.5E-3 respectively). Interestingly, AAV-Cxcl2 infected digits had significantly less bone repair with 45% decrease in new bone area and 19% decreased length as compared to AAV-GFP infection (Figure 5H, J, K; p=0.016 and 0.046, respectively). This suggests that while upregulated in blastemal fibroblasts, overexpression of Cxcl2 alone does not promote regeneration, though additional research is required to determine its exact role in digit wound healing.

**Figure 5.**
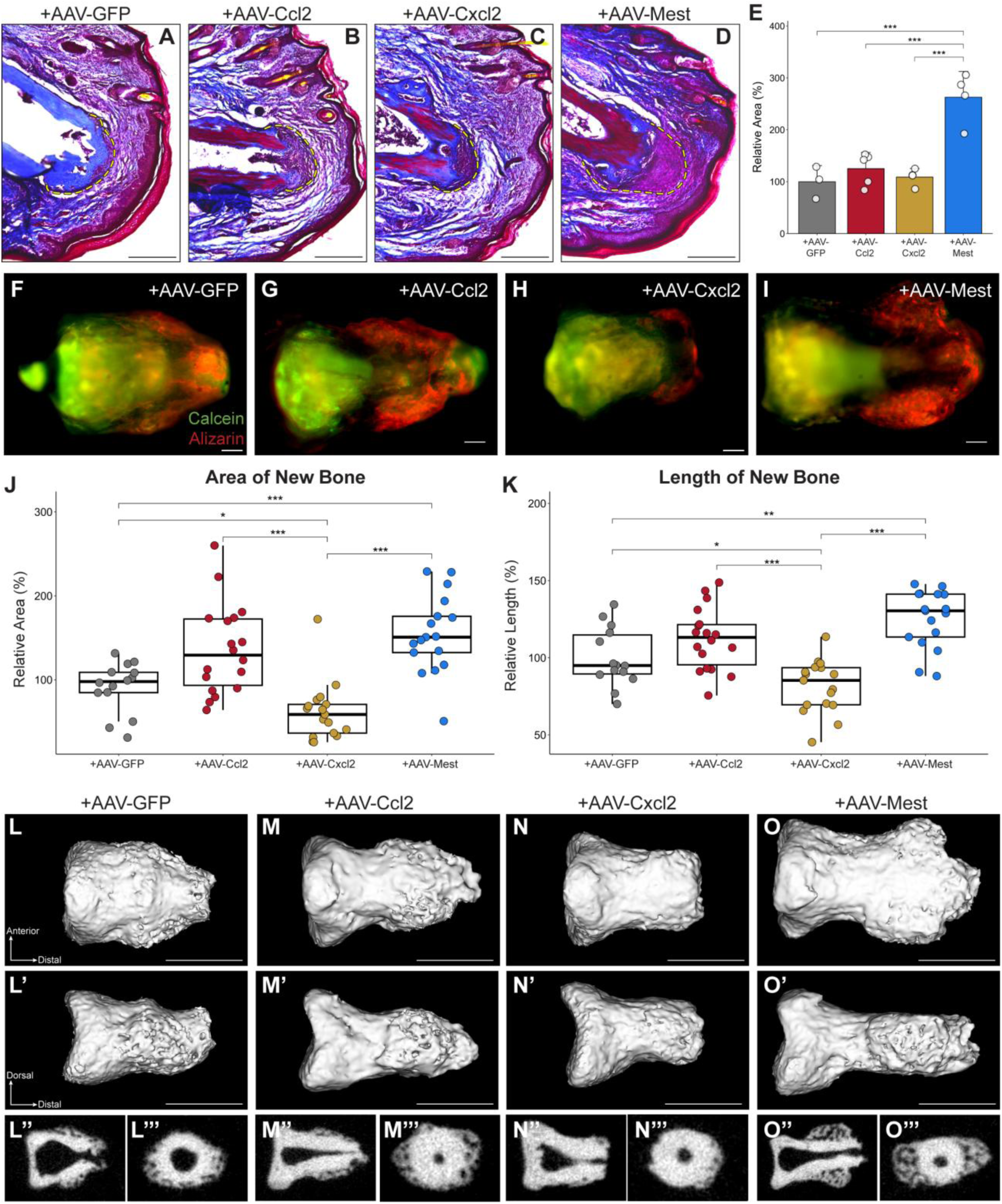
AAV delivery of candidate pro-regenerative genes induce bone growth in P2 amputation. 28dpa Masson’s trichrome stained P2 digits following AAV infection with (A) GFP, (B) Ccl2, Cxcl2, and (D) Mest; dashed yellow lines mark distal connective tissue area quantified in (E). (F-I) Representative calcein (green) and alizarin complexone (red) differentially stained P2 28dpa bones from digits infected with (F) AAV-GFP, (G) AAV-Ccl2, (H) AAV-Cxcl2, and (I) AAV-Mest. 2D quantification of new bone (red) (J) area and (K) length. Representative (L-O) dorsal and (L’-O’) lateral view microCT images of 28dpa AAV infected P2 digits. (L”-O”) and (L’’’-O’’’) are cross sections of (L-O) and (L’-O’), respectively to highlight cortical and woven bone. Scale bars in A-D = 250µm, F-I, L-O’ = 100µm. Data in (E) are reported as mean ± standard deviation and in (J, K) as median ± interquartile range; *p<0.05, **p<0.01, ***p<0.001.

We performed microCT analysis of all 28dpa P2 bones. Consistent with the calcein-alizarin differential fluorescence data, the AAV-Cxcl2 microCT scans reveal qualitatively less woven bone as compared to the AAV-GFP innately repaired bones (Figure 5L-L’’’, N-N’’’). Again, consistent with the 2D calcein-alizarin data, AAV-Ccl2 infected digits have qualitatively longer bones and additional woven bone growth (Figure M-M”’). Supporting Mest as a pro-regenerative factor, AAV-Mest infected digits had qualitatively more distal outgrowth and larger regions of woven bone deposition as compared to any of the other AAV cohorts (Figure 5O-O’’’). Axis rotation reveals the majority of this ectopic bone growth occurred along the anterior-posterior axis, resulting in a flat morphology with respect to the dorsal-ventral axis (Figure 5L’-O’), which may be due to mechanical loading during gait, though this has yet to be tested. Thus, our data indicate that while Cxcl2 impaired bone repair, expression of Ccl2 or Mest was sufficient to induce increased bone regeneration following P2 non-regenerative amputations, with Mest inducing a more dramatic phenotype.

## DISCUSSION

In this study, we aimed to identify fibroblast subpopulations and associated genes specific to P2 digit fibrosis or P3 digit regeneration. Towards this goal, we have generated two valuable resources for the field: 1) a strain and stage matched scRNAseq derived dataset through digit fibrosis and regeneration, and 2) a highly effective AAV-based local gene delivery technique to manipulate post-amputation digit gene expression in vivo. As a proof-of-concept, we focused on fibroblasts and related genes; however, the dataset and techniques utilized here are broadly applicable to the community to assess differences in fibrosis and regeneration in all cell types in the mouse digit. Our P2 digit scRNAseq analysis yielded over 12,000 fibroblasts from UA, 12, 17, and 28dpa samples (Figure 1E). Consistent with our characterization of P3 digit tip fibroblasts^6^, we also find fibroblast subtype heterogeneity among P2 derived fibroblasts with different population dynamics throughout fibrosis (Figures 1E, G). We hypothesized that there are fibroblast subtypes that are specific to P3 regeneration and/or P2 fibrosis and underlie these responses. Indeed, following computational integration we identified P2 and P3 enriched subpopulations at each matched time point (Figure 2). While the skewing of some fibroblast subtypes could be attributed to anatomical differences between the P2 and P3 digit, we experimentally limited this possibility by minimizing dissection of P3 epidermis (nail organ, sweat glands) and P2 tendons. Instead, the proportion of fibroblast subtypes residing the homeostatic P2 versus P3 digit may serve as a signature of regenerative potential (Figure 2A’, E). To explore this further, we focused on the fibroblast subtypes that were significantly skewed toward P2 fibrosis at in population size at 12dpa, 17dpa or 28dpa to identify candidate pro-fibrotic and pro-regenerative genes. Ultimately, we identified 98 putative pro-fibrotic genes and 27 putative pro-regenerative genes (Tables S4 and S6). Interestingly, the majority of candidate pro-regenerative genes, which included many of our previously reported blastemal-enriched genes^6^, are involved in the immune response. Furthermore, we found upregulation of inflammation-related pathways initially enriched in the blastema in 28dpa P2 fibroblasts, suggesting a later-stage inflammatory response in fibrosis (Figure S2E-G). This finding corroborates previous studies establishing chronic inflammation as a hallmark of fibrosis in other models^20,21^ and may prove important in developing new therapies for fibrosis.

To evaluate the biological role our computationally identified candidate genes may have, we developed and validated the usage of recombinant AAV6-based gene delivery to the post-amputation mouse P2 and P3 digits. We demonstrate that this method is ideal for local, transient gene expression in fibroblasts, with the highest expression from 3-10 days post infection (Figure 4). Though our studies were focused on infecting fibroblasts, this injection technique using different AAV pseudotypes will facilitate gene manipulation in other cell types. Additionally, timing of injection can be adjusted for desired gene overexpression temporal profile. This tool is a major step forward for unbiased functional assessment of candidate genes or other target gene manipulation in the digit fibroblasts.

In testing our computationally identified pro-fibrotic genes 28dpa AAV-Pcolce2 infected digits had longer and larger bones along the P-D axis but thinner along the A-P axis. Prelp had the reverse effect, with shorter bones that were slightly wider (Figure 4). While both are matrisome associated genes^29^, how they modulate bone shape is unknown. Prelp is expressed in myofibroblasts to increase collagen production to promote fibrosis and was shown to stiffen the ECM in zebrafish CNS wound healing, which may explain the shorter bones due to physical constraints^37,44,45^. Pcolce2 is an enzyme in procollagen cleaving also shown to have increased expression in myocardial fibrosis, though how it affected the digit bone 3D structure remains unknown^35,36^. In assessing our computationally identified pro-regenerative genes, Ccl2 and Mest overexpression resulted in increased P2 bone growth at 28dpa (Figure 5). These results complement our previous finding in which genetic knockout of Mest resulted in delayed bone regeneration^8^. Ccl2 attracts macrophages and monocytes^39^, and has been shown to promote regeneration in other models^40–42^. Whether Ccl2 and Mest are acting in the same pathway to promote bone growth is unknown. In contrast, AAV-Cxcl2 injected digits had very little bone growth, resulting in stunted bones (Figure 5). This inhibitory effect digit bone repair may be due to the role of Cxcl2 as a potent chemoattractant for neutrophils^39^, for which increased and prolonged population of has been associated with impaired wound healing^46^. However, the changes in immune cell populations with overexpression of these given genes is still unclear and would require future studies to understand their roles in digit bone repair. Additionally, to further enhance regeneration, it would be necessary to test additional genes or treat with combination of genes.

## RESOURCE AVAILABILITY

### Lead contact

Requests for further information and resources should be directed to and will be fulfilled by the lead contact, Jessica Lehoczky.

### Materials availability

All unique/stable reagents generated in this study are available from the lead contact with a completed materials transfer agreement.

### Data and code availability

- UA, 12dpa, 17dpa, and 28dpa P2 mouse digit scRNAseq data have been deposited at NCBI GEO under accession GSE293537 and are publicly available as of the date of publication.
- This paper analyzes existing, publicly available scRNAseq data accessible at NCBI GEO: GSE267446 and GSE135985.
- This paper does not report original code.
- Any additional information required to reanalyze the data reported in this paper is available from the lead contact upon request.

## ACKNOWLEDGEMENTS

We thank the Center for Cellular profiling at BWH for assistance with sample processing for 10X Genomics. We thank Dr. Julia Charles for her assistance with microCT. We acknowledge the Boston Children’s Viral Core for help with AAVs. We acknowledge the NeuroTechnology Studio at BWH for providing Zeiss LSM880 confocal microscope access. This work was supported by NICHD/NIH R01HD109200 to J.A.L. V.J received support from NIH/NIAMS T32AR055885 and F31AR082220.

## AUTHOR CONTRIBUTIONS

V.J and J.A.L conceptualized and designed the study. V.J, S.D.S, and J.S performed the experiments. V.J and J.A.L performed the computational analyses. V.J, S.D.S, and J.A.L analyzed and interpreted the data. V.J and J.A.L wrote the manuscript, and all authors approved the final version.

## SUPPLEMENTAL FIGURES

**Supplemental Figure 1.**
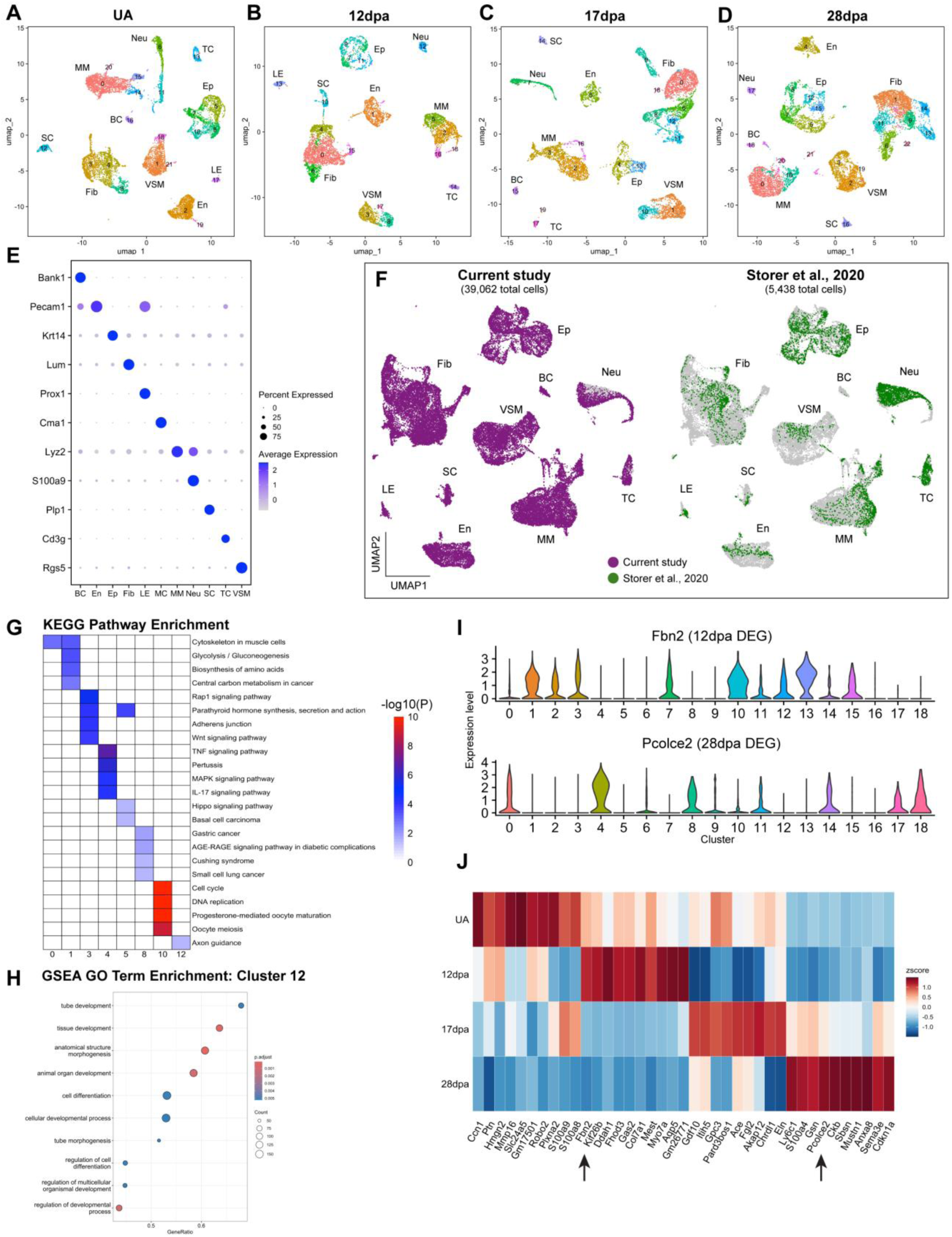
P2 individual stage scRNAseq and integrated P2 fibroblast; Related to Figure 1. UMAP plots of P2 (A) UA, (B) 12dpa, (C) 17dpa, and (D) 28dpa scRNAseq Seurat defined clusters. BC: B cells, En: endothelial cells; Ep: epithelial cells, Fib: fibroblasts, LE: lymphatic endothelial cells, MM: macrophages/monocytes, Neu: neutrophils, SC: Schwann cells, TC: T cells, VSM: vascular smooth muscle. (E) Dot plot of marker gene expression used for cell type assignment in the integrated P2 dataset (Figure 1D). (F) Validation UMAP plots of all P2 cells from this study (purple) integrated with the P2 cells from GSE135985 (green). (G) KEGG pathway enrichment for clusters 0, 1, 3, 4, 5, 8, 10, and 12 from P2 fibroblasts (Figure 1E). (H) GO term enrichment for cluster 12 from P2 fibroblasts (Figure 1E, H). (I) Top 10 DEGs for each stage of P2 fibroblasts, highlighting Fbn2 and Pcolce2 (arrows). (J) Violin plots for Fbn2 and Pcolce2 gene expression in P2 integrated fibroblast clusters.

**Supplemental Figure 2.**
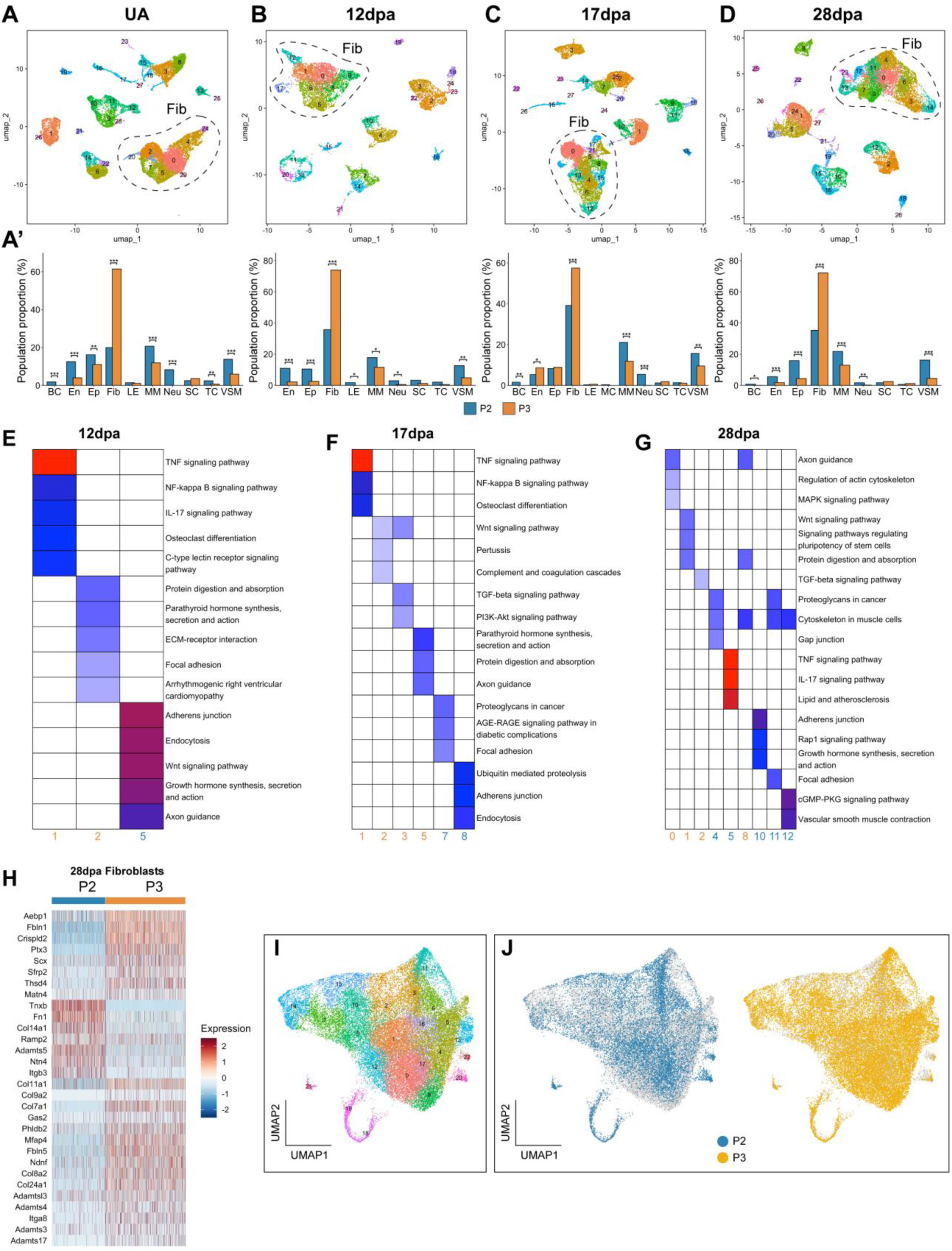
P2 and P3 stage-matched scRNAseq and fibroblast analysis; Related to Figure 2. UMAP plots of stage-matched P2 and P3 scRNAseq data for (A) UA, (B) 12dpa, (C) 17dpa, and (D) 28dpa colored by Seurat defined clusters; fibroblast populations at each time point circled with a black dashed line. (A’-D’) Differential proportion analysis of population sizes for each cell type compared between P2 and P3 digits. *p<0.05, **p<0.01, ***p<0.001. (E-G) KEGG pathway enrichment for clusters with significantly skewed population sizes between P2 and P3 fibroblasts at (E) 12dpa, (F) 17dpa, and (G) 28dpa. (H) Heatmap of DEGs associated with the ‘ECM organization’ GO term split by P2 and P3 fibroblasts at 28dpa. (I) UMAP plot of all P2 and P3 integrated fibroblasts, including 11dpa and 14dpa P3 fibroblasts; (J) colored by P2 (blue) and P3 (gold).

**Supplemental Figure 3.**
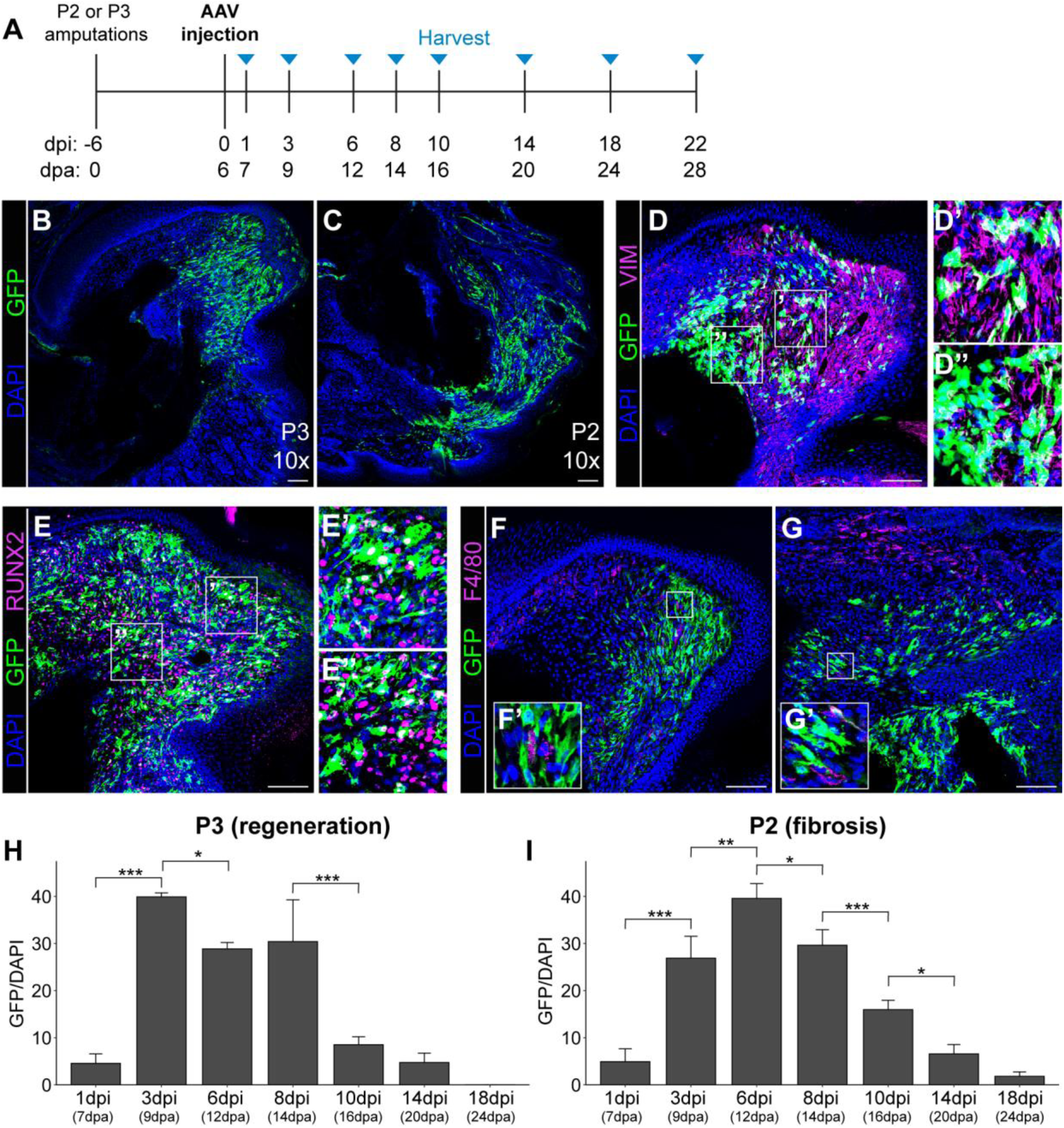
Validation of AAV2/6-GFP digit infection; Related to Figure 3, 4, 5. (A) Post-amputation digit AAV injection experimental timeline; days post injection (dpi) are 6 days offset from days post-amputation (dpa). (B, C) Anti-GFP IHC in (B) 9dpa P3 and (C) 14dpa P2 AAV-GFP infected digits. (D, E) IHC for GFP (green) and (D) VIM and (E) RUNX2 (magenta) in 16dpa P3 AAV-GFP infected digits where white cells are double labelled; D’ and D” are insets indicated on D. (F, G) IHC for GFP (green) and F4/80 (magenta) in (F) 9dpa P2 and (G) 9dpa P2 AAV-GFP infected digits; high magnification insets (F’, G’) marked by white boxes. (H, I) Image quantification of GFP signal 1-, 3-, 8-, 10-, 14- and 18dpi of (G) P3 and (H) P2 digits. Data are reported as mean ± standard deviation; p-values are only shown for comparisons of adjacent time points. *p<0.05, **p<0.01, ***p<0.001. Scale bars = 100µm.

**Supplemental Figure 4.**
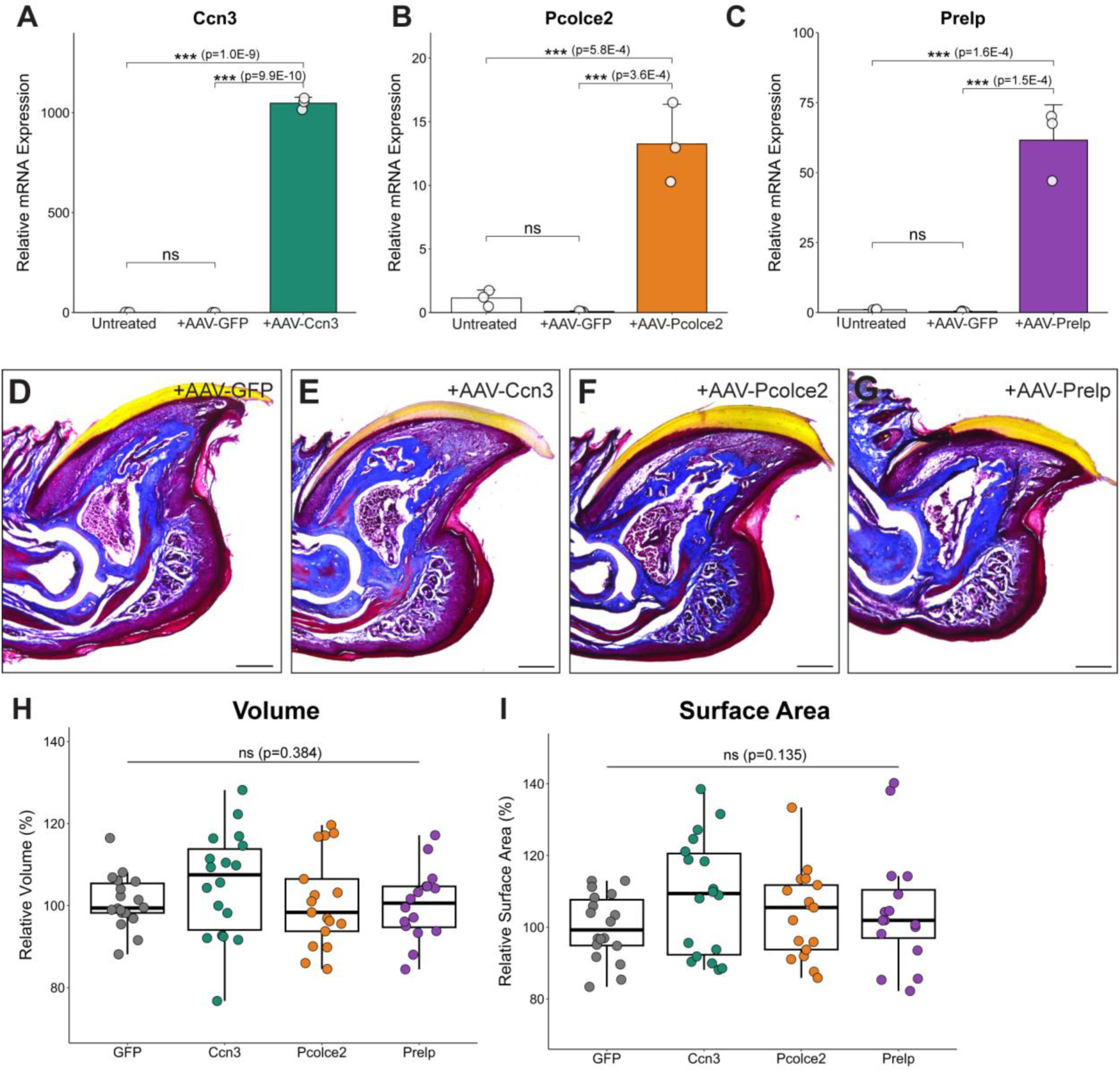
P3 AAV delivery of candidate pro-fibrotic genes; Related to Figure 4. (A-C) qPCR analysis for (A) Ccn3, (B) Pcolce2, and (C) Prelp expression in untreated and AAV-infected 12dpa P3 digits. (D-G) Masson’s trichrome stained sections of 28dpa digits following AAV infection of (D) GFP, (E) Ccn3, (F) Pcolce2, and (G) Prelp. (H-I) Quantification (H) volume and (I) surface area of 28dpa AAV-GFP, AAV-Ccn3, AAV-Pcolce2, and AAV-Prelp injected digits from P3 bone µCT analysis. Scale bars = 250µm. Data in (A-C) are reported as mean ± standard deviation and in (H, I) as median ± interquartile range. ***p<0.01, ns = not significant.

**Supplemental Figure 5.**
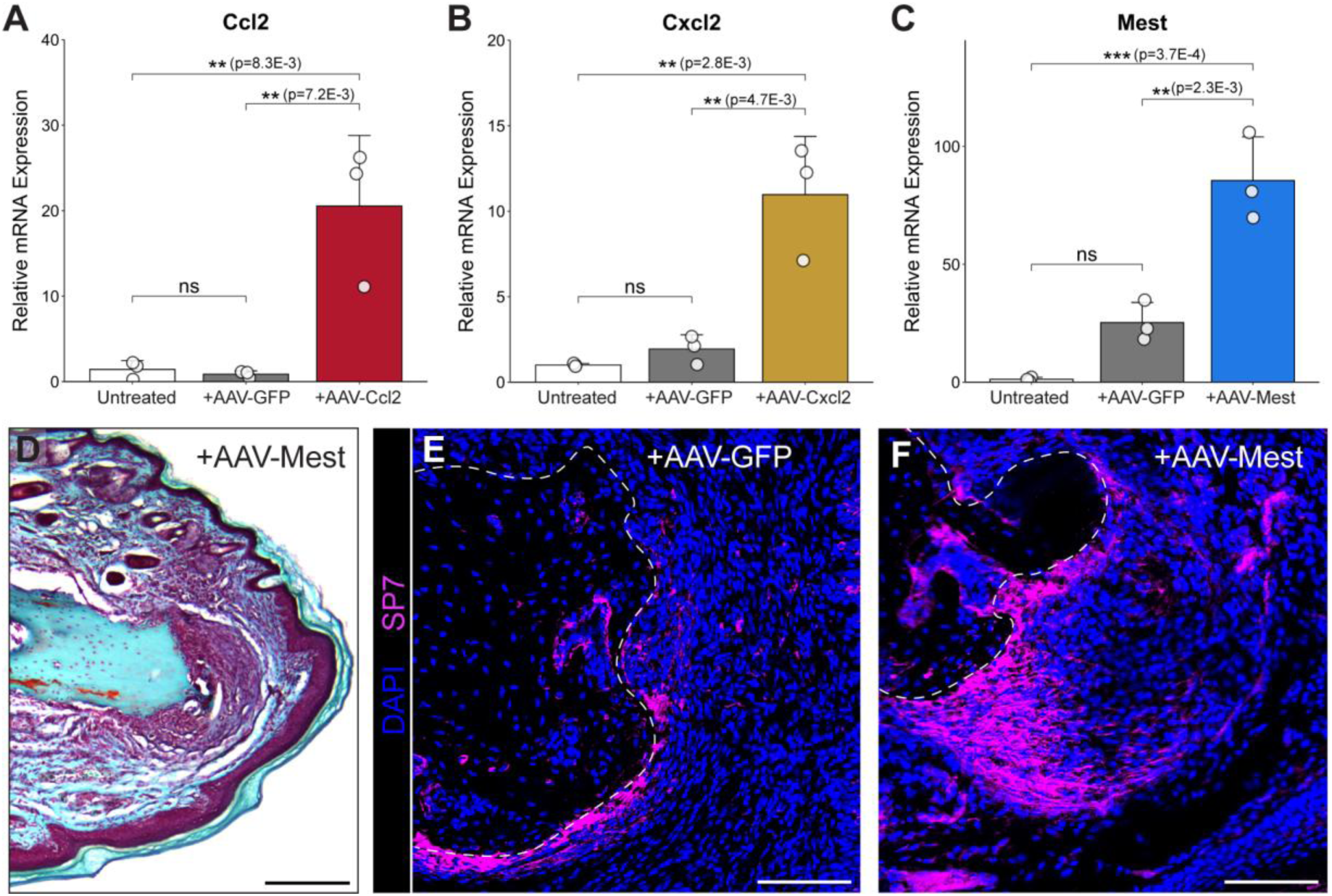
P2 AAV delivery of candidate pro-regenerative genes, Related to Figure 5. (A-C) qPCR analysis for (A) Ccl2, (B) Cxcl2, and (C) Mest expression in untreated and AAV-infected 12dpa P2 digits. (D) Safranin O/Fast Green stained sections of 28dpa digits following AAV-Mest P2 infection. (E, F) Anti-SP7 (magenta) IHC in 28dpa P2 digits following (E) AAV-GFP and (F) AAV-Mest infection. Scale bar = 250µm. Data are reported as mean ± standard deviation. **p<0.01, ***p<0.001, ns = not significant.

## SUPPLEMENTAL TABLE LEGENDS

**Table S1. Differentially expressed genes in P2 fibroblast clusters that exhibited significant population change through time; Related to Figures 1 and S1.**

Genes significantly upregulated in clusters 0, 1, 3, 5, 8, 10, and 12 of P2 fibroblasts identified by FindAllMarkers in Seurat. Genes were further filtered for avg_log2FC > 0.58, p_val_adjust < 0.05, and min.pct > 0.25. Tabs contains data for above clusters. Columns include gene by NCBI gene ID, average log fold change (avg_log2FC) calculated against all other clusters, pct.1 (percentage of cells in the relevant cluster with specific gene expression), pct.2 (percentage of cells in all other clusters with specific gene expression), adjusted p-value (p_val_adj) and cluster.

**Table S2. Differentially expressed genes for each time point in P2 fibroblasts; Related to Figure S1.**

Genes significantly upregulated in each time point (UA, 12dpa, 17dpa, and 28dpa) identified by FindMarkers in Seurat and further filtered for avg_log2FC > 0.58, p_val_adjust < 0.05, and min.pct > 0.25. Each tab contains data for distinct post-amputation time point. Columns include gene by NCBI gene ID, average log fold change (avg_log2FC) calculated against all other times, pct.1 (percentage of cells in the relevant time sample with specific gene expression), pct.2 (percentage of cells in all other time samples with specific gene expression), adjusted p-value (p_val_adj) and cluster.

**Table S3. Differentially expressed genes in significantly skewed P2 clusters at 28dpa; Related to Figures 2 and S2.**

Genes significantly upregulated in clusters 3, 4, 5, 10, 11, and 12 of 28dpa P2 and P3 integrated fibroblasts identified by FindAllMarkers in Seurat. Genes were further filtered for avg_log2FC > 0.58, p_val_adjust < 0.05, and min.pct > 0.25. Each tab contains data for above clusters. Columns include gene by NCBI gene ID, average log fold change (avg_log2FC) calculated against all other clusters, pct.1 (percentage of cells in the relevant cluster with specific gene expression), pct.2 (percentage of cells in all other clusters with specific gene expression), adjusted p-value (p_val_adj) and cluster.

**Table S4. Candidate pro-fibrotic genes; Related to Figures 2, 4, S2, and S4**

Candidate pro-fibrotic genes (NCBI gene ID) in fibroblasts as determined by identifying genes in Table S2 at 28dpa time point in common with Table S3.

**Table S5. Differentially expressed genes in significantly skewed P3 clusters at 12dpa; Related to Figures 2 and S2.**

Genes significantly upregulated in clusters 1 and 2 of 12dpa P2 and P3 integrated fibroblasts identified by FindAllMarkers in Seurat. Genes were further filtered for avg_log2FC > 0.58, p_val_adjust < 0.05, and min.pct > 0.25. Each tab contains data for above clusters. Columns include gene by NCBI gene ID, average log fold change (avg_log2FC) calculated against all other clusters, pct.1 (percentage of cells in the relevant cluster with specific gene expression), pct.2 (percentage of cells in all other clusters with specific gene expression), adjusted p-value (p_val_adj) and cluster.

**Table S6. Candidate pro-regenerative genes; Related to Figures 2, 5, S2, and S5**

Candidate pro-regenerative genes (NCBI gene ID) in fibroblasts as determined by identifying similar genes in Table S5 and previously determined blastema enriched genes^6^.

**Table S7. Primers; Related to Figures 4, 5, S4, and S5.**

All primers used in this study are listed, including primers for cloning candidate genes and primers for qPCR.

## MATERIALS AND METHODS

### Mice

All mice used in this study were housed in BWH Hale BTM vivarium maintained by the Center for Comparative Medicine. All mouse protocols and procedures were approved by the BWH IACUC. 8 to 10-week-old male and female FVB/NJ mice (JAX #001800) were used for all experiments in this study.

### Digit amputation surgery and AAV injection

For digit amputation surgeries, mice were anesthetized using inhaled 1-2% isoflurane in oxygen. Under stereomicroscope visualization, hindlimb digits 2, 3, and 4 were amputated using a sterile #11 scalpel. P3 digit tip regenerative amputations removed half of the third phalangeal segment; P2 non-regenerative digit amputations were made midway through the second phalangeal segment. Subcutaneous meloxicam (5mg/kg) and buprenorphine HCl (0.05mg/kg) were given for pre- and post-surgical analgesia.

For AAV injections, mice were anesthetized using inhaled 1-2% isoflurane and given meloxicam (5mg/kg) for analgesia. For all AAV serotypes and constructs, 2 µL of high titer (>1×10^13^ vg/mL) AAV was directly injected into each digit at 6dpa using a 5uL Hamilton syringe and 33-gauge removeable 0.5 inch needle with a 12 degree bevel. To validate pseudotype infection specificity and amount, three digits on the right hindpaw were injected with the same AAV-GFP pseudotype while the left hindpaw digits remained untouched. For all other studies, all 6 digits on each mouse were injected with the same pseudotype and recombinant AAV. Mice who received the same AAV were co-housed.

### Single-cell RNA sequencing (scRNAseq)

For each scRNAseq time point (UA, 12dpa, 17dpa, and 28dpa), stage-specific P2 digit tissue was dissected and pooled from four mice (24 digits). Dissections were performed under microscopy to minimize epidermis and tendons; UA homeostatic tissue was collected from the tissue along the P2 bone. Single-cell suspensions for each time point were generated as previously described^6^, with the addition of propidium iodide for live-cell sorting. In short, each pooled sample underwent enzymatic digestion with 2.5% trypsin and 10% collagenase followed by trituration by pipet. Red blood cells were removed with ACK lysing buffer and cell suspension was strained twice through a 35µm filter. Each sample was suspended in 0.4% BSA/PBS with 1µg/µL of propidium iodide. BD FACS Aria sorter was used to collect propidium iodide negative cells for each time point. Single-cell droplet encapsulation and library construction were performed at the BWH Center for Cellular Profiling utilizing the 10X Chromium single-cell gene expression platform (10X Genomics). Single-cell 3’ v3 chemistry (10x Genomics) chemistry was utilized to generate cDNA libraries and all libraries were sequenced by Illumina NextSeq 500 at the Dana Farber Cancer Institute Molecular Biology Core Facility.

### scRNAseq computational analyses

Computational analyses were performed using the O2 high-performance computing environment maintained by the Research Computing Group at Harvard Medical School. For each P2 dataset, CellRanger 7.1.0 (10x Genomics) was used to align raw FASTQ files to the mouse mm10-2020-A reference transcriptome, with introns. Subsequent analyses were performed with Seurat v5.1.0^47^ package for R v4.3.1^48^. Raw data contained 16045, 8813, 10514, and 15253 sequenced cells for UA, 12-, 17-, and 28dpa, respectively. Low quality cells and doublets were removed^49^. Filtered datasets were used for all downstream analyses and included: P2 samples of 11834, 7208, 8286, and 11752 cells for UA, 12-, 17-, and 28dpa time points; GSE267446 P3 samples of 11499, 6918, 3039, 5414, 7947, and 8591 cells for UA, 11-, 12-, 14-, 17-, and 28dpa time points; and for validation, GSE135985 P2 samples of 1165, 917, and 3356 cells for 10-, 10-, and 14dpa time points, respectively^6,7^. Each amputation and time point underwent individual normalization, identification of variable features, scaling, cluster identification and dimensional reduction.

Integration of datasets across amputations and/or time was performed using the CCA Integration method in Seurat followed by re-normalization, scaling, cluster identification and dimensional reduction. The broad fibroblast population for each dataset was identified and subsetted by marker genes Pdgfrα and Lumican, followed by re-scaling and clustering. Differentially expressed genes (DEGs) for subpopulations were identified using the Seurat function FindAllMarkers with min.pct = 0.25, adj.pval < 0.05, and avg.log2FC > 1.00. DEGs between time points and between P2 and P3 fibroblasts were identified using the Seurat function FindMarkers with min.pct = 0.25, adj.pval < 0.05, and avg.log2FC > 0.58. All GO term and KEGG pathway enrichment analyses were performed using ClusterProfiler v4.10.0^50^. Statistical significance for fibroblast subpopulation changes through time or between P2 and P3 contribution was determined using a previously described differential proportion analysis pipeline^51^. SeuratExtend^52^ was utilized to calculate Z-scores for DEGs and generation of Z-score heatmaps.

### Generation of recombinant AAV constructs and stocks

AAV-CAG-GFP pseudotypes (2/6, 2/8, 2/9, 2/7m8, and 2/PHP.S) were obtained from the Boston Children Hospital Viral Core. Recombinant AAV constructs were cloned into the pAAV2-CAG-tdTomato backbone (addgene #59462) where tdTomato was excised with BamHI and EcoRI restriction enzymes. Full length mouse cDNAs for Ccl2 (NM_011333.3), Ccn3 (NM_010930.5), Cxcl2 (NM_009140.2), Mest (NM_008490.2), Pcolce2 (NM_029620.2), and Prelp (NM_054077.4) were amplified from oligoDT primed mouse e12.5 cDNA libraries. Gene specific primers with AAV backbone homology tails were used for amplification with Primestar GLX DNA polymerase (Takara Bio) (Table S7). Amplicons were used for Gibson assembly using NEB HiFi Assembly Mix and clones were maintained in NEB Stable E. coli (New England Biolabs). All plasmids were DNA sequence verified by whole plasmid sequencing at Plasmidsaurus. Endotoxin-free plasmid DNA for each construct was prepared and SignaGen Laboratories for AAV6 production. All AAV stocks received were verified to be of viral titer > 1×10^13^ vg/mL.

### Section immunohistochemistry (IHC) and histology

Collected digits were fixed in 4% PFA at 4° C overnight, followed by decalcification with Decalcifying Solution-Lite (Sigma-Aldrich) or with Surgipath Decalcifier I (Leica Biosystems). Tissues were processed through a 5% to 30% sucrose gradient, then embedded in OCT freezing medium (Tissue-Tek) for cryosectioning on a Leica CM3050s cryostat.

For IHC, 18µm sections were blocked with 10% goat serum in PBST and incubated with primary antibody at 4° C overnight; anti-GFP (1:1000; Aves Labs GFP-1020), anti-VIM (1:200, Novus NB300-233), anti-RUNX2 (1:200; Abcam ab192256), anti-F4/80 (1:100; BioRad MCA497GA), anti-SP7 (1:2000, Abcam ab22552). Tissues were incubated with appropriate Cy3 or A647 conjugated secondary antibodies for one hour at room temperature followed by PBST washes and DAPI staining. Slides were mounted in Fluoromount (Southern Biotech) and imaged on a Zeiss LSM880 confocal microscope, capturing z-stack images. Max intensity projections were generated and processed in Fiji^53^. GFP per DAPI signal values for each relevant image were quantified in Fiji, with DAPI quantified utilizing the thresholding and analyze particles functions with follow-up manual annotations. Three areas with high GFP expression for each representative image were chosen for quantification. Statistical significance was assessed by one-way ANOVA, followed by post-hoc t-tests with Bonferroni correction for multiple hypotheses.

14µm sections were made for 28dpa histological staining. Masson’s trichrome Staining Kit (Polysciences) was used for trichrome staining following the manufacturer’s protocol for frozen sections. For Safranin O/Fast Green staining, slides were stained with Safranin O and Fast Green (Sigma-Aldrich) as previously described^54^. Brightfield imaging was performed with a Leica DM2000 upright microscope followed by image processing and analysis in Fiji. The area of connective tissue aggregation distal to the P2 bone was calculated in Fiji and statistical significance was assessed by one-way ANOVA followed by post-hoc t-tests with Bonferroni correction.

### Quantitative PCR (qPCR)

To validate overexpression of each gene of interest following AAV injection, 18 P2 or P3 injected digits were collected at 12dpa (6dpi). Tissues were homogenized and lysed in TRIzol (Invitrogen) using Navy Eppendorf RNA lysis tubes in a Bullet Blender Storm24 machine (Next Advance) at 4° C. Subsequent RNA extraction was performed following standard TRIzol protocol. cDNA was generated utilizing the SuperScript IV First Strand Synthesis kit primed with oligo-dT (Invitrogen). qPCR was performed with gene specific primers for Ccl2, Ccn3, Cxcl2, Mest, Pcolce2, Prelp (Table S7) including those previously published^34,45,55–58^ and SsoAdvanced Universal SYBR green mastermix (BioRad) on a QuantStudio 5 Real-time PCR machine. ΔCt was calculated using Gapdh as a housekeeping gene with in-plate technical triplicates. Relative mRNA expression was calculated using the 2^(-ΔΔCt)^ method with normalization to the respective untreated sample. All qPCR experiments were further performed in triplicate. Statistical significance was evaluated by one-way ANOVA followed by post-hoc t-test with Bonferroni correction.

### Whole mount digit bone analysis

Alizarin red skeletal staining was used to assess P3 bone regeneration, 18 digits per AAV group were collected at 28dpa and stained as previously reported^59^. All stained P3 bones were imaged in 50% glycerol with a Leica M165FC stereomicroscope. Quantification of P3 2D area and bone length was performed in Fiji. Bone length was measured from the midpoint of the P2/P3 joint to the distal tip of the bone. Statistical significance was calculated by one-way ANOVA followed by post-hoc t-tests with Bonferroni correction for multiple hypothesis testing.

Differential calcein and alizarin complexone staining was used to assess new bone growth post P2 amputation. For all AAV groups, 10mg/kg intraperitoneal calcein (Sigma-Aldrich) was given immediately following P2 amputations and 30mg/kg intraperitoneal alizarin complexone (Sigma-Aldrich) was given two days prior (26dpa) to harvest^60^. 18 digits per AAV group were collected at 28dpa and dehydrated in 100% EtOH at 96° C for 24 hours. Digits were cleared in 1% KOH for 24 hours and then stored in 50% glycerol. Digits were imaged with a Leica M165FC epifluorescent steromicroscope. Images were processed and new bone growth was quantified in Fiji. Pre-existing bone was masked and remaining Alizarin-stained bone was quantified to determine new bone growth area and length of the bone from amputation site utilizing the distal Calcein front to the distal tip of bone. For the untreated compared to AAV-GFP P2 bones, statistical significance was determined using two-sample t-tests. For the AAV candidate gene P2 bones, statistical significance was calculated using one-way ANOVA followed by post-hoc t-test with Bonferroni correction.

All P3 alizarin red and P2 calcein-alizarin complexone 28dpa digits also underwent microCT scanning. Digits were scanned on a Scanco Medical µCT 35 system at 55kVp, 145µA, 8W with 600ms integration time and voxel size of 7µm in 25-50% glycerol. Each scan was converted to a DICOM and analyzed using the 3D Slicer image computing platform^61^. 3D segmentation models were made for both P2 and P3 scanned bones; volume and surface area were quantified for the P3 bones. Statistical significance of 3D volume and surface area was calculated by one-way ANOVA followed by post-hoc t-tests with Bonferroni correction. To evaluate dorsal-ventral (D-V) and anterior-posterior (A-P) P3 bone widths while controlling for length, the proximal-distal (P-D) length of each digit was scaled to 0-100%. The mean and standard deviation for D-V and A-P bone widths was calculated at every 2% of P-D length. Statistical significance of D-V and A-P bone widths were assessed by MANOVA followed by one-way ANOVA and post-hoc t-tests with Benjamini-Hochberg correction at each interval measured along the P-D length. Statistically significant regions for indication in Figure 4L was determined by a minimum of two consecutive intervals each with P < 0.05.

## REFERENCES

1. Borgens, R.B. (1982). Mice regrow the tips of their foretoes. Science (80-.). 10.1126/science.7100922.

2. Neufeld, D.A., and Zhao, W. (1995). Bone regrowth after digit tip amputation in mice is equivalent in adults and neonates. Wound Repair Regen. 3, 461–466. 10.1046/j.1524-475X.1995.30410.x.

3. Fernando, W.A., Leininger, E., Simkin, J., Li, N., Malcom, C.A., Sathyamoorthi, S., Han, M., and Muneoka, K. (2011). Wound healing and blastema formation in regenerating digit tips of adult mice. Dev. Biol. 350, 301–310. 10.1016/J.YDBIO.2010.11.035.

4. Rinkevich, Y., Lindau, P., Ueno, H., Longaker, M.T., and Weissman, I.L. (2011). Germ-layer and lineage-restricted stem/progenitors regenerate the mouse digit tip. Nature. 10.1038/nature10346.

5. Lehoczky, J.A., Robert, B., and Tabin, C.J. (2011). Mouse digit tip regeneration is mediated by fate-restricted progenitor cells. Proc. Natl. Acad. Sci. U. S. A. 108, 20609– 20614. 10.1073/pnas.1118017108.

6. Johnson, G.L., Masias, E.J., and Lehoczky, J.A. (2020). Cellular Heterogeneity and Lineage Restriction during Mouse Digit Tip Regeneration at Single-Cell Resolution. Dev. Cell. 10.1016/j.devcel.2020.01.026.

7. Storer, M.A., Mahmud, N., Karamboulas, K., Borrett, M.J., Yuzwa, S.A., Gont, A., Androschuk, A., Sefton, M. V., Kaplan, D.R., and Miller, F.D. (2020). Acquisition of a Unique Mesenchymal Precursor-like Blastema State Underlies Successful Adult Mammalian Digit Tip Regeneration. Dev. Cell 52, 509–524.e9. 10.1016/j.devcel.2019.12.004.

8. Jou, V., Peña, S., and Lehoczky, J.A. (2024). Regeneration-specific promoter switching facilitates Mest expression in the mouse digit tip to modulate neutrophil response. npj Regen. Med., 1–49. 10.1038/s41536-024-00376-w.

9. Neufeld, D.A., and Zhao, W. (1993). Phalangeal regrowth in rodents: postamputational bone regrowth depends upon the level of amputation. Prog. Clin. Biol. Res.

10. Schotté, O.E., and Smith, C.B. (1959). Wound Healing Processes in Amputated Mouse Digits. Biol. Bull. 117, 546–561.

11. Jou, V., and Lehoczky, J.A. (2023). Toeing the line between regeneration and fibrosis. Front. Cell Dev. Biol. 11, 1–12. 10.3389/fcell.2023.1217185.

12. Yu, L., Dawson, L.A., Yan, M., Zimmel, K., Lin, Y.L., Dolan, C.P., Han, M., and Muneoka, K. (2019). BMP9 stimulates joint regeneration at digit amputation wounds in mice. Nat. Commun. 10, 1–9. 10.1038/s41467-018-08278-4.

13. Wu, Y., Wang, K., Karapetyan, A., Fernando, W.A., Simkin, J., Han, M., Rugg, E.L., and Muneoka, K. (2013). Connective Tissue Fibroblast Properties Are Position-Dependent during Mouse Digit Tip Regeneration. PLoS One 8. 10.1371/journal.pone.0054764.

14. Shyh-Chang, N., Zhu, H., Yvanka De Soysa, T., Shinoda, G., Seligson, M.T., Tsanov, K.M., Nguyen, L., Asara, J.M., Cantley, L.C., and Daley, G.Q. (2013). Lin28 enhances tissue repair by reprogramming cellular metabolism. Cell 155, 778. 10.1016/j.cell.2013.09.059.

15. Yu, L., Han, M., Yan, M., Lee, J., and Muneoka, K. (2012). BMP2 induces segment-specific skeletal regeneration from digit and limb amputations by establishing a new endochondral ossification center. Dev. Biol. 372, 263–273. 10.1016/j.ydbio.2012.09.021.

16. Yu, L., Dawson, L.A., Yan, M., Zimmel, K., Lin, Y.-L., Dolan, C.P., Han, M., and Muneoka, K. (2019). BMP9 stimulates joint regeneration at digit amputation wounds in mice. Nat. Commun. 2019 101 10, 1–9. 10.1038/s41467-018-08278-4.

17. Lee, J., Marrero, L., Yu, L., Dawson, L.A., Muneoka, K., and Han, M. (2013). SDF-1α/CXCR4 signaling mediates digit tip regeneration promoted by BMP-2. Dev. Biol. 382, 98–109. 10.1016/j.ydbio.2013.07.020.

18. Agrawal, V., Kelly, J., Tottey, S., Daly, K.A., Johnson, S.A., Siu, B.F., Reing, J., and Badylak, S.F. (2011). An isolated cryptic peptide influences osteogenesis and bone remodeling in an adult mammalian model of digit amputation. Tissue Eng Part A 17, 3033–3044. 10.1089/ten.TEA.2011.0257.

19. Mu, X., Bellayr, I., Pan, H., Choi, Y., and Li, Y. (2013). Regeneration of soft tissues is promoted by MMP1 treatment after digit amputation in mice. PLoS One 8, e59105. 10.1371/journal.pone.0059105.

20. Wynn, T.A., and Ramalingam, T.R. (2012). Mechanisms of fibrosis: Therapeutic translation for fibrotic disease. Nat. Med. 18, 1028–1040. 10.1038/nm.2807.

21. Antar, S.A., Ashour, N.A., Marawan, M.E., and Al-Karmalawy, A.A. (2023). Fibrosis: Types, Effects, Markers, Mechanisms for Disease Progression, and Its Relation with Oxidative Stress, Immunity, and Inflammation. Int. J. Mol. Sci. 24. 10.3390/ijms24044004.

22. Lu, Y. (2004). Recombinant Adeno-Associated Virus As Delivery Vector for Gene Therapy - A Review. Stem Cells Dev. 13, 133–145. 10.1089/154732804773099335.

23. Naso, M.F., Tomkowicz, B., Perry, W.L., and Strohl, W.R. (2017). Adeno-Associated Virus (AAV) as a Vector for Gene Therapy. BioDrugs 31, 317–334. 10.1007/s40259-017-0234-5.

24. Wang, D., Tai, P.W.L., and Gao, G. (2019). Adeno-associated virus vector as a platform for gene therapy delivery. Nat. Rev. Drug Discov. 18, 358–378. 10.1038/s41573-019-0012-9.

25. Westhaus, A., Cabanes-Creus, M., Rybicki, A., Baltazar, G., Navarro, R.G., Zhu, E., Drouyer, M., Knight, M., Albu, R.F., Ng, B.H., et al. (2020). High-Throughput In Vitro, Ex Vivo, and In Vivo Screen of Adeno-Associated Virus Vectors Based on Physical and Functional Transduction. Hum. Gene Ther. 31. 10.1089/hum.2019.264.

26. Piras, B.A., Tian, Y., Xu, Y., Thomas, N.A., O’Connor, D.M., and French, B.A. (2016). Systemic injection of AAV9 carrying a periostin promoter targets gene expression to a myofibroblast-like lineage in mouse hearts after reperfused myocardial infarction. Gene Ther. 23, 469–478. 10.1038/gt.2016.20.

27. Sharma, A., Ghosh, A., Hansen, E.T., Newman, J.M., and Mohan, R.R. (2010). Transduction efficiency of AAV 2/6, 2/8 and 2/9 vectors for delivering genes in human corneal fibroblasts. Brain Res. Bull. 81, 273–278. 10.1016/j.brainresbull.2009.07.005.

28. Shwartz, Y., Gonzalez-Celeiro, M., Chen, C.L., Pasolli, H.A., Sheu, S.H., Fan, S.M.Y., Shamsi, F., Assaad, S., Lin, E.T.Y., Zhang, B., et al. (2020). Cell Types Promoting Goosebumps Form a Niche to Regulate Hair Follicle Stem Cells. Cell 182, 578–593.e19. 10.1016/j.cell.2020.06.031.

29. Hynes, R.O., and Naba, A. (2012). Overview of the matrisome-An inventory of extracellular matrix constituents and functions. Cold Spring Harb. Perspect. Biol. 4, 1–16. 10.1101/cshperspect.a004903.

30. Yin, H., Liu, N., Zhou, X., Chen, J., and Duan, L. (2023). The advance of CCN3 in fibrosis. J. Cell Commun. Signal. 17, 1219–1227. 10.1007/s12079-023-00778-3.

31. Marchal, P.O., Kavvadas, P., Abed, A., Kazazian, C., Authier, F., Koseki, H., Hiraoka, S., Boffa, J.J., Martinerie, C., and Chadjichristos, C.E. (2015). Reduced NOV/CCN3 expression limits inflammation and interstitial renal fibrosis after obstructive nephropathy in mice. PLoS One 10, 1–12. 10.1371/journal.pone.0137876.

32. Ren, Z., Hou, Y., Ma, S., Tao, Y., Li, J., Cao, H., and Ji, L. (2014). Effects of CCN3 on fibroblast proliferation, apoptosis and extracellular matrix production. Int. J. Mol. Med. 33, 1607–1612. 10.3892/ijmm.2014.1735.

33. Riser, B.L., Najmabadi, F., Perbal, B., Peterson, D.R., Rambow, J.A., Riser, M.L., Sukowski, E., Yeger, H., and Riser, S.C. (2009). CCN3 (NOV) is a negative regulator of CCN2 (CTGF) and a novel endogenous inhibitor of the fibrotic pathway in an in vitro model of renal disease. Am. J. Pathol. 174, 1725–1734. 10.2353/ajpath.2009.080241.

34. Betageri, K.R., Link, P.A., Haak, A.J., Ligresti, G., Tschumperlin, D.J., and Caporarello, N. (2023). The matricellular protein CCN3 supports lung endothelial homeostasis and function. Am. J. Physiol. - Lung Cell. Mol. Physiol. 324, L154–L168. 10.1152/ajplung.00248.2022.

35. Thomas, M.J., Xu, H., Wang, A., Beg, M.A., and Sorci-Thomas, M.G. (2024). PCPE2: Expression of multifunctional extracellular glycoprotein associated with diverse cellular functions. J. Lipid Res. 65, 100664. 10.1016/j.jlr.2024.100664.

36. Baicu, C.F., Zhang, Y., Van Laer, A.O., Renaud, L., Zile, M.R., and Bradshaw, A.D. (2012). Effects of the absence of procollagen C-endopeptidase enhancer-2 on myocardial collagen accumulation in chronic pressure overload. Am. J. Physiol. - Hear. Circ. Physiol. 303, 234–240. 10.1152/ajpheart.00227.2012.

37. Zhang, Y., Fu, C., Zhao, S., Jiang, H., Li, W., and Liu, X. (2022). PRELP promotes myocardial fibrosis and ventricular remodelling after acute myocardial infarction by the wnt/β–catenin signalling pathway. Cardiovasc. J. Afr. 33.

38. Ding, F., Zheng, P., Yan, X. yue, Chen, H. jian, Fang, H. ting, Luo, Y. yuan, Peng, Y. xuan, Zhang, L., and Yan, Y. e. (2024). Adipocyte-secreted PRELP promotes adipocyte differentiation and adipose tissue fibrosis by binding with p75NTR to activate FAK/MAPK signaling. Int. J. Biol. Macromol. 279, 135376. 10.1016/j.ijbiomac.2024.135376.

39. Sokol, C.L., and Luster, A.D. (2015). The chemokine system in innate immunity. Cold Spring Harb. Perspect. Biol. 7, 1–20. 10.1101/cshperspect.a016303.

40. Wang, W., Chen, X.K., Zhou, L., Wang, F., He, Y.J., Lu, B.J., Hu, Z.G., Li, Z.X., Xia, X.W., Wang, W.E., et al. (2024). Chemokine CCL2 promotes cardiac regeneration and repair in myocardial infarction mice via activation of the JNK/STAT3 axis. Acta Pharmacol. Sin. 45, 728–737. 10.1038/s41401-023-01198-0.

41. Jablonski, C.L., Leonard, C., Salo, P., and Krawetz, R.J. (2019). CCL2 But Not CCR2 Is Required for Spontaneous Articular Cartilage Regeneration Post-Injury. J. Orthop. Res. 37, 2561–2574. 10.1002/jor.24444.

42. Toya, M., Zhang, N., Tsubosaka, M., Kushioka, J., Gao, Q., Li, X., Chow, S.K.H., and Goodman, S.B. (2023). CCL2 promotes osteogenesis by facilitating macrophage migration during acute inflammation. Front. Cell Dev. Biol. 11, 1–10. 10.3389/fcell.2023.1213641.

43. Cheng, N., Kim, K.H., and Lau, L.F. (2022). Senescent hepatic stellate cells promote liver regeneration through IL-6 and ligands of CXCR2. JCI Insight 7. 10.1172/jci.insight.158207.

44. Kolb, J., Tsata, V., John, N., Kim, K., Möckel, C., Rosso, G., Kurbel, V., Parmar, A., Sharma, G., Karandasheva, K., et al. (2023). Small leucine-rich proteoglycans inhibit CNS regeneration by modifying the structural and mechanical properties of the lesion environment. Nat. Commun. 14. 10.1038/s41467-023-42339-7.

45. Yamauchi, Y., Mieno, H., Suetsugu, H., Watanabe, H., and Nakaya, M. (2024). Elevated PRELP expression in heart and liver fibrosis promotes collagen production. Biochem. Biophys. Res. Commun. 734, 150785. 10.1016/j.bbrc.2024.150785.

46. Wilgus, T.A., Roy, S., and McDaniel, J.C. (2013). Neutrophils and Wound Repair: Positive Actions and Negative Reactions. Adv. Wound Care 2, 379–388. 10.1089/wound.2012.0383.

47. Stuart, T., Butler, A., Hoffman, P., Hafemeister, C., Papalexi, E., Mauck, W.M. 3rd, Hao, Y., Stoeckius, M., Smibert, P., and Satija, R. (2019). Comprehensive Integration of Single-Cell Data. Cell 177, 1888–1902.e21. 10.1016/j.cell.2019.05.031.

48. R Core Team (2018). R: A Language and Environment for Statistical Computing.

49. McGinnis, C.S., Murrow, L.M., and Gartner, Z.J. (2019). DoubletFinder: Doublet Detection in Single-Cell RNA Sequencing Data Using Artificial Nearest Neighbors. Cell Syst. 8, 329–337.e4. 10.1016/j.cels.2019.03.003.

50. Yu, G., Wang, L.G., Han, Y., and He, Q.Y. (2012). ClusterProfiler: An R package for comparing biological themes among gene clusters. Omi. A J. Integr. Biol. 16, 284–287. 10.1089/omi.2011.0118.

51. Farbehi, N., Patrick, R., Dorison, A., Xaymardan, M., Janbandhu, V., Wystub-Lis, K., Ho, J.W.K., Nordon, R.E., and Harvey, R.P. (2019). Single-cell expression profiling reveals dynamic flux of cardiac stromal, vascular and immune cells in health and injury. Elife. 10.7554/eLife.43882.

52. Hua, Y., Weng, L., Zhao, F., Rambow, F., and Essen, H. (2024). SeuratExtend : Streamlining Single-Cell RNA-Seq Analysis Through an Integrated and Intuitive Framework. bioRxiv.

53. Schindelin, J., Arganda-Carreras, I., Frise, E., Kaynig, V., Longair, M., Pietzsch, T., Preibisch, S., Rueden, C., Saalfeld, S., Schmid, B., et al. (2012). Fiji: an open-source platform for biological-image analysis. Nat. Methods 9, 676–682. 10.1038/nmeth.2019.

54. Leite, C.B.G., Fricke, H.P., Tavares, L.P., Nshimiyimana, R., Mekhail, J., Kilgallen, E., Killick, F., Whalen, J.D., Lehoczky, J.A., Serhan, C.N., et al. (2025). Maresin 1-LGR6 axis mitigates inflammation and posttraumatic osteoarthritis after transection of the anterior cruciate ligament in mice. Osteoarthr. Cartil., 1–13. 10.1016/j.joca.2025.03.005.

55. Pollard, R.D., Blesso, C.N., Zabalawi, M., Fulp, B., Gerelus, M., Zhu, X., Lyons, E.W., Nuradin, N., Francone, O.L., Li, X.A., et al. (2015). Procollagen C-endopeptidase enhancer protein 2 (PCPE2) reduces atherosclerosis in mice by enhancing scavenger receptor class B1 (SR-BI)-mediated high-density lipoprotein (HDL)-cholesteryl ester uptake. J. Biol. Chem. 290, 15496–15511. 10.1074/jbc.M115.646240.

56. Nakawaki, M., Uchida, K., Miyagi, M., Inoue, G., Kawakubo, A., Kuroda, A., Satoh, M., and Takaso, M. (2020). Sequential CCL2 Expression Profile After Disc Injury in Mice. J. Orthop. Res. 38, 895–901. 10.1002/jor.24522.

57. Sano, M., Ijichi, H., Takahashi, R., Miyabayashi, K., Fujiwara, H., Yamada, T., Kato, H., Nakatsuka, T., Tanaka, Y., Tateishi, K., et al. (2019). Blocking CXCLs–CXCR2 axis in tumor–stromal interactions contributes to survival in a mouse model of pancreatic ductal adenocarcinoma through reduced cell invasion/migration and a shift of immune-inflammatory microenvironment. Oncogenesis 8. 10.1038/s41389-018-0117-8.

58. Anunciado-Koza, R.P., Manuel, J., Mynatt, R.L., Zhang, J., Kozak, L.P., and Koza, R.A. (2017). Diet-induced adipose tissue expansion is mitigated in mice with a targeted inactivation of mesoderm specific transcript (Mest). PLoS One 12, 1–29. 10.1371/journal.pone.0179879.

59. Johnson, G.L., Glasser, M.B., Charles, J.F., Duryea, J., and Lehoczky, J.A. (2022). En1 and Lmx1b do not recapitulate embryonic dorsal-ventral limb patterning functions during mouse digit tip regeneration. Cell Rep. 41, 111701. 10.1016/j.celrep.2022.111701.

60. Castilla-Ibeas, A., Zdral, S., Galán, L., Haro, E., Allou, L., Campa, V.M., Icardo, J.M., Mundlos, S., Oberg, K.C., and Ros, M.A. (2023). Failure of digit tip regeneration in the absence of Lmx1b suggests Lmx1b functions disparate from dorsoventral polarity. Cell Rep. 42. 10.1016/j.celrep.2022.111975.

61. Fedorov, A., Beichel, R., Kalpathy-Cramer, J., Finet, J., Fillion-Robin, J.C., Pujol, S., Bauer, C., Jennings, D., Fennessy, F., Sonka, M., et al. (2012). 3D Slicer as an image computing platform for the Quantitative Imaging Network. Magn. Reson. Imaging 30, 1323–1341. 10.1016/j.mri.2012.05.001.

